# The molecular and cellular interplay between the osteopetrosis-associated proteins SNX10, OSTM1, and CLC-7 during osteoclastogenesis

**DOI:** 10.1101/2025.03.31.646258

**Authors:** Nina Reuven, Sabina Winograd-Katz, Maayan Barnea-Zohar, Amir Pri-Or, Yishai Levin, Jean Vacher, Benjamin Geiger, Ari Elson

## Abstract

Bone-resorbing osteoclasts (OCLs) are large, multi-nucleated cells that are formed through well-regulated differentiation and cell fusion of monocyte-macrophage precursors. Disruption of OCL-mediated bone resorption perturbs bone formation, remodeling, and homeostasis that, in turn, can lead to severe illnesses, such as autosomal recessive osteopetrosis (ARO). Mutations in the intracellular trafficking-associated protein sorting nexin 10 (SNX10) lead to “OCL-rich” ARO, in which OCLs are produced but are inactive. Furthermore, OCL fusion is deregulated in SNX10-knockout (SKO) mice: mature mutant OCLs fuse continuously to generate gigantic cells, in vitro and in vivo, unlike wild-type OCLs that stop fusing with each other upon maturation. Mutations in CLC-7, the lysosomal voltage-gated Cl^-^/H^+^ exchanger, and OSTM1, the beta-chain of the exchanger, also induce ARO in humans and in mouse models, and are associated with the presence of large OCLs. This study explored the molecular interplay between SNX10, CLC-7 and OSTM1 by directly comparing the phenotypes of cultured OCLs lacking one of these proteins. We show that loss of SNX10, OSTM1, or CLC-7 leads to the formation of similarly-gigantic OCLs in culture, due to deregulated fusion between mature OCLs that proceeds with similar kinetics. All three proteins are associated with LAMP1-positive lysosomes, localized in both perinuclear and peripheral regions of mature wild-type OCLs. Co-immunoprecipitation studies indicated that SNX10 physically interacts with CLC-7. Notably, SNX10-KO OCLs exhibited a significant reduction in peripheral lysosomes containing CLC-7 and OSTM1, suggesting that SNX10 is required for their transport to the cell periphery. Taken together, these findings indicate that SNX10 regulates the subcellular distribution of lysosomes containing CLC-7 and OSTM1, thereby controlling both the fusion and functionality of mature OCLs.

## Introduction

Osteoclasts (OCLs) are large, multi-nucleated phagocytes that degrade bone matrix and play a key role in bone homeostasis and remodeling [1–3]. These cells originate from monocyte/macrophage precursor cells through differentiation and cell-cell fusion that are driven by the cytokines M-CSF and RANKL [4, 5]. Mature, functional OCLs adhere tightly to bone and secrete protons and proteases that together degrade the mineral and organic components of the underlying bone matrix. Secretion occurs through the directed fusion of lysosomes with the plasma membrane at a specialized, resorption-associated region of the ventral OCL membrane known as the ruffled border. This region is surrounded by a belt of podosomal adhesions, termed the Sealing Zone Structure in OCLs grown on resorbable surface, which physically delimits and isolates the bone surface area undergoing degradation [2, 6, 7].

Bone degradation by OCLs is tightly regulated and is typically balanced by bone production by osteoblasts, to maintain bone mass and structural integrity. Upsetting this physiological balance can cause serious skeletal diseases, as occurs, for example, in bone-invading cancer, where excessive OCL activity enhances survival of bone metastasis, and in osteoporosis, where the imbalanced action of osteoblasts and OCLs leads to pathological reductions in bone mass [7, 8]. Conversely, insufficient bone resorption can also lead to serious pathological states. A severe example is autosomal recessive osteopetrosis (ARO, also known as infantile malignant osteopetrosis), which occurs at an incidence of 1:250,000 live births in the general population [9, 10]. ARO is characterized by, among other symptoms, massively increased bone mass, pathological bone fractures, growth retardation, and bone marrow failure. ARO presents shortly after birth and is typically fatal during the first decade of life although, notably, in some cases hematopoietic stem cell transplantation can halt disease progression and even reverse some of its symptoms [9–12].

The genetic basis for ARO is heterogeneous. “OCL-poor” ARO, which is characterized by the absence of OCLs, is caused by inactivating mutations in genes that are essential for osteoclastogenesis, such as *TNFSF11* and *TNFRSF11A*, which encode the key osteoclastogenic cytokine RANKL and its receptor RANK, respectively [10]. “OCL-rich” ARO, characterized by the presence of inactive OCLs, is caused by mutations that disrupt resorption-related cellular functions [10]. Approximately 50% of cases of OCL-rich ARO are caused by mutations in the *TCIRG1* gene, which encodes the A3 subunit of the V0 complex of the vacuolar H^+^-ATPase responsible for acidifying the bone-OCL interface [9, 13, 14]. A further 23% of cases result from mutations in *CLCN7*, which encodes the CLC-7 protein of the voltage-gated Cl^-^/H^+^ exchanger that participates in the acidification of the ‘resorption lacuna’ [15–18], or in *OSTM1*, its β subunit [9, 19–21]. Another 5% of ARO cases are linked to mutations in *SNX10*, encoding sorting nexin 10, a protein involved in vesicular trafficking [22], while isolated cases have been attributed to mutations in other genes, such as *PLEKHM1* [9]. Despite the involvement of these proteins in ARO, the interplay between them in wild-type (WT) and in osteopetrotic OCLs is still poorly understood.

Mouse models that recapitulate major ARO symptoms provide valuable insights into disease mechanisms. For example, mice lacking CLC-7 [18, 23, 24], OSTM1 [21, 25, 26], or SNX10 [27–29] are massively osteopetrotic, exhibit severe developmental defects, and survive poorly. Some ARO-associated mutations also affect tissues beyond bone, as demonstrated by the severe neurological deficiencies observed both in humans suffering from mutations in CLC-7 or OSTM1 and in the corresponding mouse models [15, 20, 30–32]. We previously demonstrated that OCLs cultured from SNX10-knockout (SKO) mice or from those homozygous for the ARO-inducing mutation R51Q SNX10 (RQ mice) are gigantic due to continuous fusion between pairs of mature OCLs [29, 33]. In contrast, fusion between mature WT OCLs is extremely rare, resulting a population of cells with a defined and reproducible size range [29, 33]. A similar situation occurs also *in vivo*, when endogenous GFP-tagged OCLs growing inside tibias of RQ or SKO mice are visualized by advanced 3D microscopy [29]. These mutant cells fuse excessively to generate OCLs that are significantly larger than their WT counterparts, and their growth is often limited by the physical constraints of the surrounding bone matrix. Collectively, these findings indicate that the size and resorption capacity of OCLs are determined by a fusion-arresting mechanism that is active, genetically-regulated, OCL cell-autonomous, and SNX10-dependent. Initial studies suggest that the hyper-fusion phenotype of the RQ OCLs may result from impaired endocytosis, which leads to the abnormal retention of fusion-promoting proteins at the plasma membrane [29, 33].

The similarities in the bone-specific symptoms induced by SNX10, OSTM1, and CLC-7 mutations in humans and in mice raise important mechanistic questions regarding the role of SNX10 in OCL fusion and resorption, and its potential interaction with OSTM1 and CLC-7. Previous studies revealed that OCLs cultured from OSTM1-knockout (OKO) mice are larger than WT controls due to excessive fusion [25], and large OCLs were also observed in the residual bone marrow space of CLC-7-knockout (CKO) mice [23]. However, the effects of loss of CLC-7, OSTM1, or SNX10 have not been compared directly in OCLs, particularly concerning the regulation of cell fusion and cell size.

In this study, we demonstrate that cultured OCLs deficient in SNX10, OSTM1, or CLC-7 all exhibit excessive and persistent fusion, resulting in the formation of gigantic OCLs. This fusion occurs between large, mature OCLs with similar kinetics across all three mutant types. Additionally, CLC-7, OSTM1, and SNX10 localize to punctate LAMP1-containing lysosomal structures distributed both at the periphery and perinuclear regions of WT OCLs, with SNX10 partially co-localizing with CLC-7 and OSTM1at both locations. Furthermore, SNX10 and CLC-7 co-immunoprecipitate, indicating that they are present in the same molecular complex. Notably, loss of SNX10 significantly reduces CLC-7- and OSTM1- containing lysosomes located at the cell periphery while leaving their perinuclear localization unaffected, suggesting that SNX10 regulates lysosomal transport to the cell surface. Collectively, these findings indicate that SNX10 acts upstream of CLC-7 and OSTM1, and that these three proteins function jointly within the same mechanism that regulates the development, size, and resorptive activity of OCLs.

## Materials and Methods

### Growth and lentiviral transduction of RAW264.7 cells

RAW264.7 cells were obtained from the ATCC. RAW264.7 cells expressing pBABEpuro-LifeAct-EGFP were described [33]. The cells were grown in Dulbecco’s Modified Eagle’s Medium (DMEM, Sigma-Aldrich, St Louis, MO, USA), supplemented with 10% fetal calf serum, 2 mM glutamine, 50 units/ml penicillin, and 50 μg/ml streptomycin. RAW 264.7 cells were grown on plastic tissue culture plates and induced to differentiate with 20 ng/ml M-CSF (Peprotech, Rehovot, Israel) and 20 ng/ml RANKL (R&D Systems, Minneapolis, MN, USA). The lentivirus expression vector pLenti-UbC-blast, which confers blasticidin resistance, was constructed by combining the backbone of pLenti6 (ThermoFisher) with a fragment encoding the ubiquitin C (UbC) promoter taken from pUltra-hot (gift from Malcolm Moore, Addgene plasmid # 24130). Mouse *Ostm1* cDNA [34] carrying a C-terminal V5 tag was subcloned into this vector by PCR. Mouse *Clcn7* cDNA was PCR-amplified from RAW264.7 OCL cDNA, and cloned into the pLenti-UbC-blast vector; all DNAs amplified by PCR were verified by sequencing. Lentivirus transducing particles were prepared in HEK293T cells as described [35]. RAW264.7 LifeAct-GFP cells were transduced for 8-16h, followed by three washes with PBS. Cells were returned to medium without selection for 8-16h, followed by selection using 2.5-5μg/ml blasticidin (BIOPrep, Omer, Israel)

### Knocking out *Snx10*, *Ostm1*, and *Clcn7* in RAW264.7 cells

RAW264.7 *Snx10* knockout cells have been described [33]. CRISPR-mediated gene knockout of *Clcn7* and *Ostm1* was performed in LifeAct-expressing RAW264.7. Cells were transfected with a plasmid providing blasticidin resistance, along with pU6-(BsaI)_CBh-Cas9-T2A- mCherry (a gift from Prof. Yosef Shaul, Weizmann Institute, Addgene, <u>135012</u>) expressing Cas9 and the appropriate sgRNA. The sgRNA sequences used are provided in Supplementary

Fig. 1. Cells were transfected using JetPEI (Polyplus Transfection, Illkirch, France) according to the manufacturer’s instructions. Two days post-transfection, cells were treated with 10 µg/ml blasticidin for 24 h to select for transfected cells, which were then plated as single cells in 96-well plates. Genomic DNA from individual clones was analyzed by sequencing PCR products with the targeted site at the center, generated using the primer sets in Supplementary Table 1. Sequencing results were deconvoluted using the Synthego ICE tool (https://icethermo.synthego.com/).

**Figure 1:**
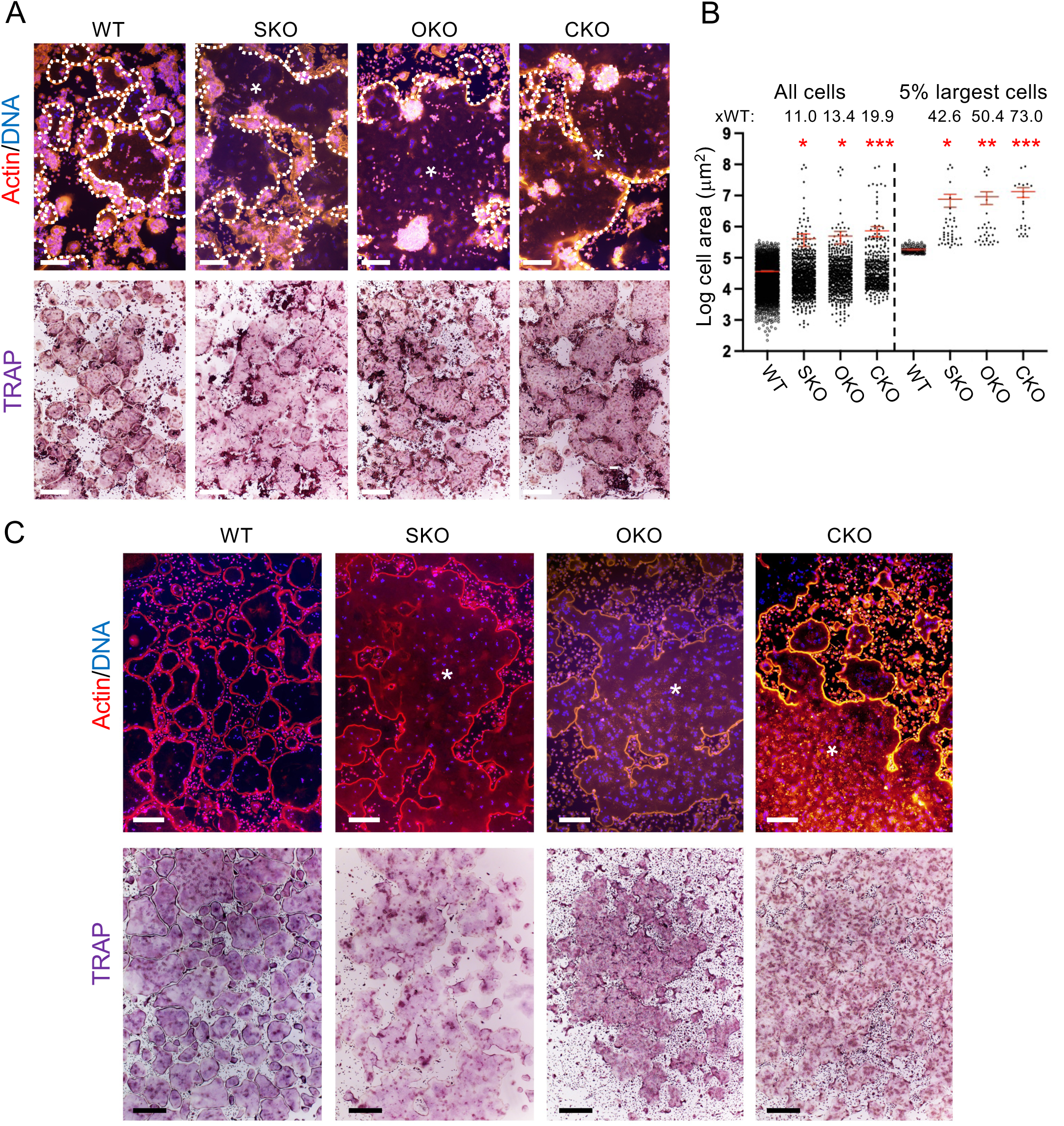
OCLs lacking SNX10 (SKO), OSTM1 (OKO), or CLCN7 (CKO) are gigantic. **A.** RAW 264.7 cells of the indicated genotypes were plated on glass coverslips and induced to differentiate with M-CSF and RANKL. Cells were stained for actin (red/orange) and DNA (blue) (top) or tartrate-resistant acid phosphatase (TRAP) (bottom). Top: Cell limits are indicated by dashed lines; the same image wihout the dashed lines in shown in Supplementary Fig. 3. Each of the mutant images contains part of a single cell (marked by an asterisk) that extends significantly beyond the field of view. Scale bars: 200 μm (top), 500 μm (bottom). WT- control, non-targeted cells. **B.** Area distribution of spleen-derived OCLs grown on glass coverslips. Left plot; The areas of multinucleated (≥three nuclei) OCLs from the indicated genotypes (OCL numbers: N=2517 WT, 831 (SKO), 606 (OKO), and 471 (CKO). Right plot: The same data for the 5% largest OCLs in each genotype. *p≤0.0244, **p=0.0073 *** p=0.0002 by one-way ANOVA, relative to WT in each graph. Also shown are the fold area increase vs. the corresponding WT cells (xWT). Red bars: Mean±SE. Note that the Y axis scale is logarithmic. **C**. Splenocytes from from mice of the indicated genotypes were grown and analyzed as in A. Asterisks in top panels indicate giant cells in the mutant cultures. Scale bars: 200 μm (top), 500 μm (bottom).

### Knock-in of an HA tag at the C-terminus of endogenous SNX10 in RAW 264.7 cells

CRISPR was used to knock in an HA tag at the C-terminus of the endogenous *Snx10* gene in LifeAct-EGFP- expressing RAW264.7 cells. To improve the yield of knock-in cells, we used a Cas9 construct that included a recruiting domain for the endogenous Mre11/Rad50/Nbs1 DNA repair complex that is needed for homology-directed repair (pU6-(BsaI)_CBh-UN- Cas9-T2A-mCherry, Addgene 135013, gift of Prof. Yosef Shaul). The donor plasmid, constructed in a pBluescript KS (Stratagene, La Jolla, CA) backbone, and the sgRNA used, are described in Supplementary Fig. 2. The blasticidin resistance cassette was amplified from a pLenti6 vector (Invitrogen/Thermo-Fisher Scientific, Waltham, MA, USA), and the homology arms were amplified from RAW264.7 genomic DNA. NEB HiFi DNA Assembly Master Mix (New England Biolabs, Ipswich, MA) was used to assemble the fragments and construct the plasmid. Cells were transfected with the Cas9/sgRNA and donor plasmids using JetPEI (Polyplus Transfection), and were selected with 2.5μg/ml blasticidin for two days, followed by 4 days growth without blasticidin. Surviving cells were plated in 96 well plates as single cells, and clones arising were analyzed by PCR of their genomic DNA. Sequencing of the PCR products verified the proper integration of the cassette, and protein blotting confirmed the expression of the SNX10-HA protein (Supplementary Fig. 2).

### Genetically-modified mice

The production, genotyping, and main phenotypes of knockout mice lacking SNX10 [29], OSTM1 [25], and CLC-7 [18] have been described. All mouse experiments described in this study were approved by the Weizmann Institute IACUC and were conducted in accordance with Israeli law.

### Culture of primary mouse OCLs from spleens

Splenocytes were used in this study for OCL formation since the massive osteopetrosis of the mutant mice prevented isolation of sufficient numbers of bone marrow cells. Spleens from 4- 8-week old mice were dissociated into PBS. Following lysis of erythrocytes, cells were seeded at a density of 2×10^6^ cells/well (WT) and 1×10^6^ cells/well (mutants) in 24-well plates, or 8×10^6^ cells/well (WT) and 4×10^6^ cells/well (mutants) in six-well plates. The cells were cultured in OCL medium (α-MEM (Sigma) supplemented with 10% fetal calf serum, 2 mM glutamine, 50 units/ml penicillin and 50 μg/ml streptomycin) supplemented with 20 ng/ml M-CSF. RANKL (20 ng/ml) was added 24-48 hours later, and the cells were grown at 37°C in 5% CO_2_ for 5-7 days with daily changes of medium. On occasion, cells were seeded onto glass coverslips inserted into 24-well plates.

### Growth of primary OCLs on fragments of bovine bone and pit resorption assay

2-3 sterile fragments (approx. 2×1×1mm) of bovine bone were placed in wells of a 24-well plate and incubated for 2-3 hours in OCL medium (lacking cytokines). Primary splenocytes were then added to the wells and grown as described above; medium was replaced every 24- 48 hours for up to 15 days. Visualization of resorption pits was performed by removing the cells by sonication in PBS followed by incubation for 2 hours in 2 μg/ml horseradish peroxidase-labelled wheat germ agglutinin (Vector Laboratories Inc, Burlingame, CA, USA), rinsing in PBS, and incubation in a solution of 3-3’-diaminobenzidine (Vector-DAB; Vector Laboratories) according to the manufacturer’s instructions.

### OCL immunofluorescence and TRAP staining

Cells were grown on glass coverslips or bone fragments. For actin staining, cells were fixed in 3% paraformaldehyde (PFA) for 20 minutes and then permeabilized in 0.5% Triton X-100/ 3% PFA for 3 min, followed by three washes in PBS. Cells were then incubated with fluorescently-conjugated phalloidin (TRITC-phalloidin, Sigma-Aldrich, P1951, phalloidin-Alexa 488, Cell Signaling Technology, Danvers, MA, USA, 8878S, or FITC-phalloidin, Sigma-Aldrich P2582, diluted 1:300-1:500 in PBS), and/or with primary antibodies for 1 hour at room temperature in PBS containing 2.5 % goat serum, followed by three washes in PBS and incubation with secondary antibodies as required. Images were captured using Olympus IX53 (Olympus, Japan) or Zeiss Axio Observer.Z1/7 (Zeiss, Germany) microscopes). The Zeiss Plan-Apochromat 63x/1.4 oil objective was used for the higher-resolution images. Ten Z-stack images (each slice 0.2μ thick) were photographed, and the ZEN 3.3 (ZEN Pro, Zeiss) software default deconvolution setting was used to improve image quality. One Z-slice was presented for each image, with the same slice shown for all channels.

Primary antibodies used for immunofluorescence were: anti-Flag-M2 (mouse monoclonal Sigma-Aldrich F1804, diluted 1:200), anti-V5 (mouse monoclonal, ThermoFisher R960 diluted 1:300, or rabbit monoclonal clone 1K20-L2, Sigma-Aldrich ZRB1509, diluted 1:300), anti-HA (mouse monoclonal clone HA.11, Babco/Covance 16B12, diluted 1:300, or rat monoclonal clone 3F10, Roche (Basel, Switzerland), diluted 1:300), anti-LAMP1 (rat monoclonal clone H4A3, Santa Cruz Biotechnology (Dallas,TX, USA) SC-20011, diluted 1:300) and anti-CLC7 (affinity-purified polyclonal antibody, diluted 1:200 [18]). Secondary antibodies, used at 1:300-1:400 dilution, were: Donkey anti-Mouse Cy5 (715-176-150), Goat anti-Mouse AF594 (115-585-166), Goat anti-Rabbit RRX (111-295-144), and Goat anti-Rat AF594 (112-585-003), all from Jackson Immunoresearch Laboratories (West Grove, PA, USA). DNA was visualized with Hoechst 33342 (Molecular Probes, Eugene, OR, USA, H- 3570, diluted 1:7000 in PBS).

Cells grown on optical plastic, glass coverslips, or bone fragments were stained for tartrate-resistant acid phosphatase (TRAP) using a Leukocyte Acid Phosphatase kit (Sigma-Aldrich). For measurement of OCL sizes in culture, primary splenocytes in 24-well plates were differentiated with M-CSF and RANKL until most of the well area contained mature, differentiated OCLs. Cells were stained for actin and DNA, photographed, and individual OCLs (≥3 nuclei/cell) were manually marked and their areas were calculated using Fiji [36]. All OCLs in an entire well or in half of it were measured.

### Phase-contrast live cell imaging and determination of “touch-to-fusion” time

Live cell imaging was performed as described previously [33]. The time between first contact between pairs of cells and their actual fusion was determined by running phase-contrast movies of cells undergoing osteoclastogenesis, such as Movies 1-4, “backwards” until we identified pairs of cells that had just fused. We then followed these cells further backwards in time until they made first contact. Mature WT OCLs did not fuse, hence in these cases we followed cell pairs from the time of first contact until they separated or until the movie ended.

### qPCR analysis

Primary OCLs were prepared from spleen cells as described above. RNA preparation and qPCR analysis were performed as in [33], with readings normalized to mouse *Rpl4*. Primers (Supplementary Table 2) were designed using Primer Express v.3 (Applied Biosystems) and validated by a standard curve and dissociation curve of the products. The fold change in target gene expression was calculated by the 2^−ΔΔCt^ relative quantification method [37] (Applied Biosystems).

### Immunoprecipitation and protein blot analysis

Cells were lysed in NP40 buffer (50mM Tris-Cl pH 7.5, 150mM NaCl, 1%NP40) supplemented with protease inhibitor cocktail (Sigma 8340 or equivalent) for 20min on ice, followed by centrifugation at 16,000 x *g* at 4°C. For immunopreciptiation, cell extracts (1.5 mg protein/RAW264.7 cells or 0.5 mg protein/HEK293T cells) were incubated with mouse anti-Flag M2 agarose (Sigma A2220), mouse anti-HA agarose (Sigma A2095), or mouse anti-V5 agarose (Sigma A7345) beads for 2-4h at 4°C, with rotation, followed by four washes with the cell lysis buffer. Proteins were eluted by incubating with 0.5-1mg/ml Flag or HA peptide, for 10min at 37°C. SDS-PAGE and protein blotting were performed as described [38]. Primary antibodies used for protein blotting were anti-FLAG (mouse monoclonal clone M2, Sigma-Aldrich F3165, diluted 1:2000), anti-β actin (mouse monoclonal clone AC-15, Sigma-Aldrich A1978, diluted 1:5000), anti-SNX10 (rabbit polyclonal, Sigma-Aldrich SAB2107086, diluted 1:1000), and the antibodies described above for immunofluorescence, diluted as follows: anti-V5 (1:1000), anti-HA (mouse monoclonal, 1:5000; rat monoclonal 1:2000), anti-CLC7 (1:1000). Secondary horseradish peroxidase-labeled goat-anti-mouse IgG (115-035-146), goat anti-rat light chain IgG (112-035-175), and goat-anti-rabbit IgG (111-035-144) antibodies were obtained from Jackson Immunoresearch Laboratories. Enhanced chemiluminescence was visualized using an Imagequant LAS 4000 Mini instrument (GE Healthcare Biosciences, Uppsala, Sweden).

### Proteomics

#### Sample Preparation

Samples were lysed using 5% SDS, reduced, alkylated, and digested with trypsin (enzyme-to-protein ratio 1:50) on S-Trap microcolumns (Protifi, NY, USA) according to the manufacturer’s instructions, as described in [39].

#### Liquid Chromatography and Mass Spectrometry

Peptide separation and mass spectrometric analysis were performed as in [1] using split-less nano-Ultra Performance Liquid Chromatography (nanoAcquity; Waters, MA, USA; 10 kpsi) coupled to a quadrupole Orbitrap mass spectrometer (Q Exactive HF; Thermo Scientific, MA, USA) via the Proxeon FlexIon nanospray apparatus, with minor modifications - The LC gradient and MS acquisition mode were adjusted as detailed below.

Peptides were eluted using a mobile phase that was composed of: A) H_2_O + 0.1% formic acid and B) acetonitrile + 0.1% formic acid, with the following gradients: 4% to 33% solvent B over 105 min, 33% to 90% B over 5 min, held at 90% B for 5 min, and then returned to 4% B. Targeted analysis was conducted in parallel reaction monitoring (PRM) mode. Ions were initially acquired at a resolution of 120,000 (at m/z 200) over a mass range of m/z 300–1650, with an AGC target (maximal number collected) of 3×10^6^ ions and a maximum injection time of 60 ms. Selected ions derived from the targeted proteins were then analyzed further at a resolution of 30,000 with a quadrupole isolation window of 1.7 m/z, an AGC target of 2×10^5^ ions, and a maximum injection time of 60 ms. The list of targeted peptides is shown in Supplemental Table 3.

#### Data Processing

Raw data were processed using Thermo Scientific Proteome Discoverer v2.4 (employing SequestHT and MS Amanda) against a mouse protein database with common contaminants included, and further analyzed using Skyline v20.1.

### Statistical analysis

Data were analyzed by one-way ANOVA or by the Kruskall-Wallis test, as indicated in the figure legends, with statistical significance set at p=0.05.

## Results

### Cultured osteoclasts lacking either SNX10, OSTM1, or CLC-7 are gigantic due to deregulated cell fusion

Mutations in SNX10, OSTM1, or CLC-7 lead to the formation of resorption-inactive OCLs that underlie severe “OCL rich” osteopetrosis in humans and in genetically-manipulated mice [9, 18, 21, 23, 25–29]. To directly compare the cellular phenotypes of OCLs lacking these genes, we disrupted the *Snx10*, *Ostm1*, or *Clcn7* genes in RAW264.7 cells (Supplementary Fig. 1). When cultured on plastic or glass surfaces and induced to differentiate with M-CSF and RANKL, non-targeted WT cells fused and developed into mature OCLs of variable sizes. In contrast, cultures of OCLs that lacked OSTM1, CLC-7, or SNX10 were dominated by much larger and rapidly-expanding OCLs (Fig. 1A upper panel, Supplementary Fig. 3). These WT and all mutant OCLs stained positively for TRAP, and possessed a peripheral podosomal array that is typical for OCLs growing on non-resorbable surfaces (Fig. 1A).

Similar findings were observed in primary OCLs derived from splenocytes of mice lacking SNX10 (SKO mice), OSTM1 (OKO mice) or CLC-7 (CKO mice) that were treated with M-CSF and RANKL (Fig. 1C, upper panel). Upon differentiation, WT control cultures contained large numbers of mature, juxtaposed OCLs with projected areas that rarely exceeded 3×10^5^ μm^2^ (Fig. 1B). While similar mature OCLs were found also in the three mutant cultures, these cultures were dominated by much larger mature OCLs, whose projected areas often reached 1×10^8^ μm^2^ (Fig. 1B). All primary OCLs in this study contained a single peripheral podosome array and stained positively for TRAP, although TRAP staining was less intense in the mutant OCLs than in the WT controls (Fig. 1C, bottom panel).

To examine the morphology of mutant OCLs grown on physiological, resorbable surfaces, primary splenocytes were seeded on fragments of bovine bone, differentiated into OCLs, and stained for actin/DNA or TRAP. Cells from SKO, OKO, and CKO mice formed gigantic OCLs that were considerably larger than those derived from WT mice (Fig. 2A). The podosomes of WT OCLs were arranged in one or several internal Sealing Zone rings per cell, which is characteristic of OCLs growing on bone surfaces (Fig. 2B). In contrast, all mutant OCLs formed a single peripheral podosome belt per cell (Fig. 2B), an organization that is typical of OCLs growing on non-resorbable surfaces. While TRAP staining was present in mutant OCLs, it was less intense than in WT OCLs (Fig. 2A). Consistent with previous studies [18, 25, 29], OCLs from three mutant strains were inactive and failed to from resorption pits when grown on bone (Supplementary Fig. 4).

**Figure 2:**
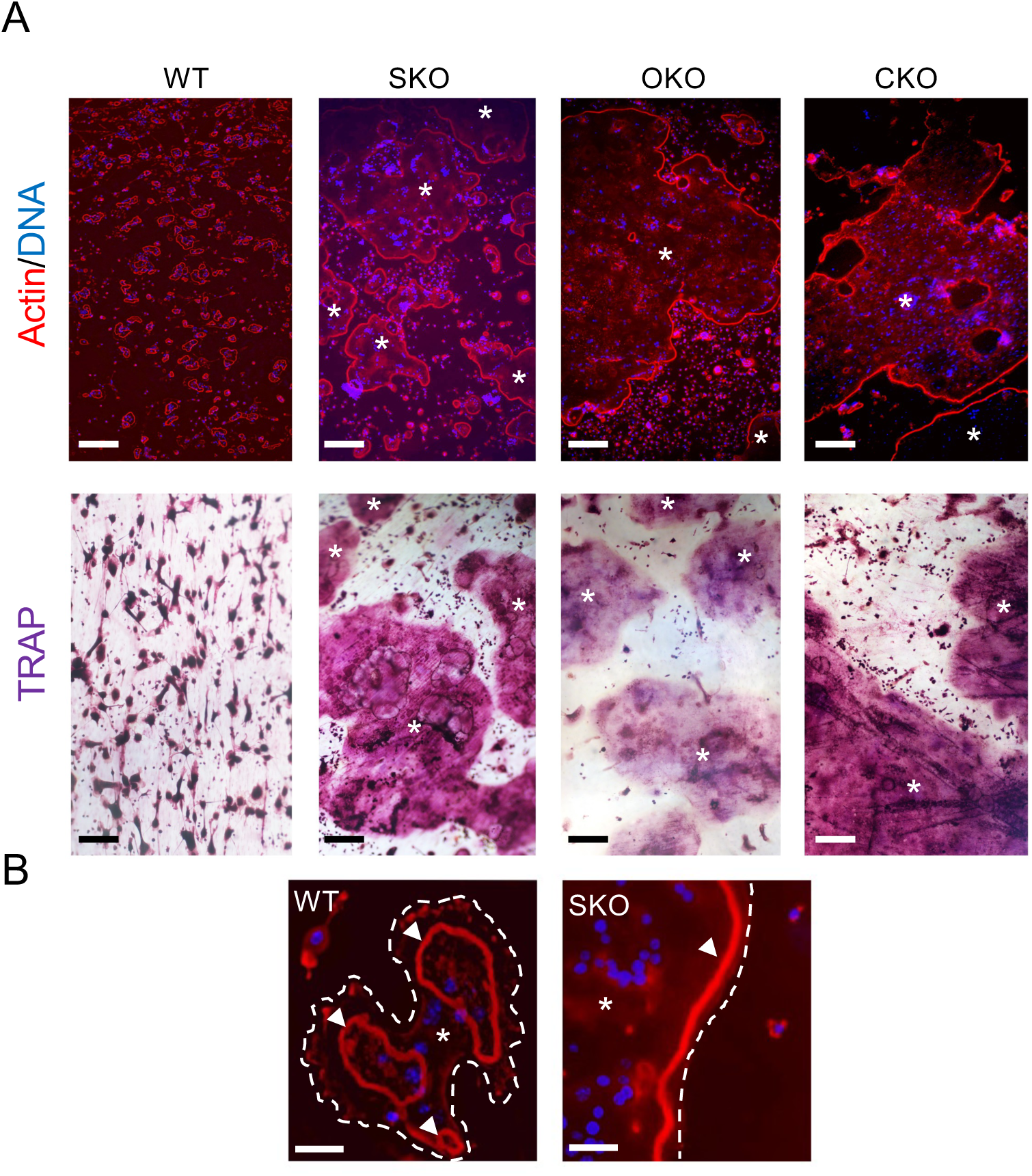
Primary OCLs deficient for SNX10, OSTM1, or CLC-7 are gigantic when cultured on bone. **A.** Primary splenocytes from mice of the indicated genotypes were seeded on fragments of bovine bone, induced to differentiate with M-CSF and RANKL, and stained (top) for actin (red) and DNA (blue) or (bottom) TRAP. Asterisks in mutant images mark cell bodies; most mutant cells are too large to fit into the field shown. Scale bars: 200 μm. **B**. Higher-power images of WT (left) and SKO (right) OCLs grown on bone, stained for actin/DNA. Dashed lines indicate cell boundaries, asterisks indicate cell bodies, arrowheads indicate podosomal SZs. Scale bars: 25 μm.

### Giant OCLs lacking SNX10, OSTM1, or CLC-7 arise from a similarly deregulated hyper-fusion process

To examine the cellular dynamics that underlie the formation of giant OCLs by the three mutant strains, we used microscopy-based live cell imaging. As previously demonstrated [33], WT mononucleated cells exposed to RANKL ceased proliferation, fused, and developed into mature, large and circular OCLs (Movie 1). Notably, these mature WT OCLs did not fuse with other mature OCLs, although they continued to fuse with immature, mono- or oligo-nucleated cells. Fusion ceased altogether when mature OCLs became completely surrounded by other mature OCLs, with which they did not fuse (Fig. 3, Movie 1). Consequently, differentiated cultures of primary WT OCLs contained large areas of individual juxtaposed mature OCLs of varying sizes.

**Figure 3:**
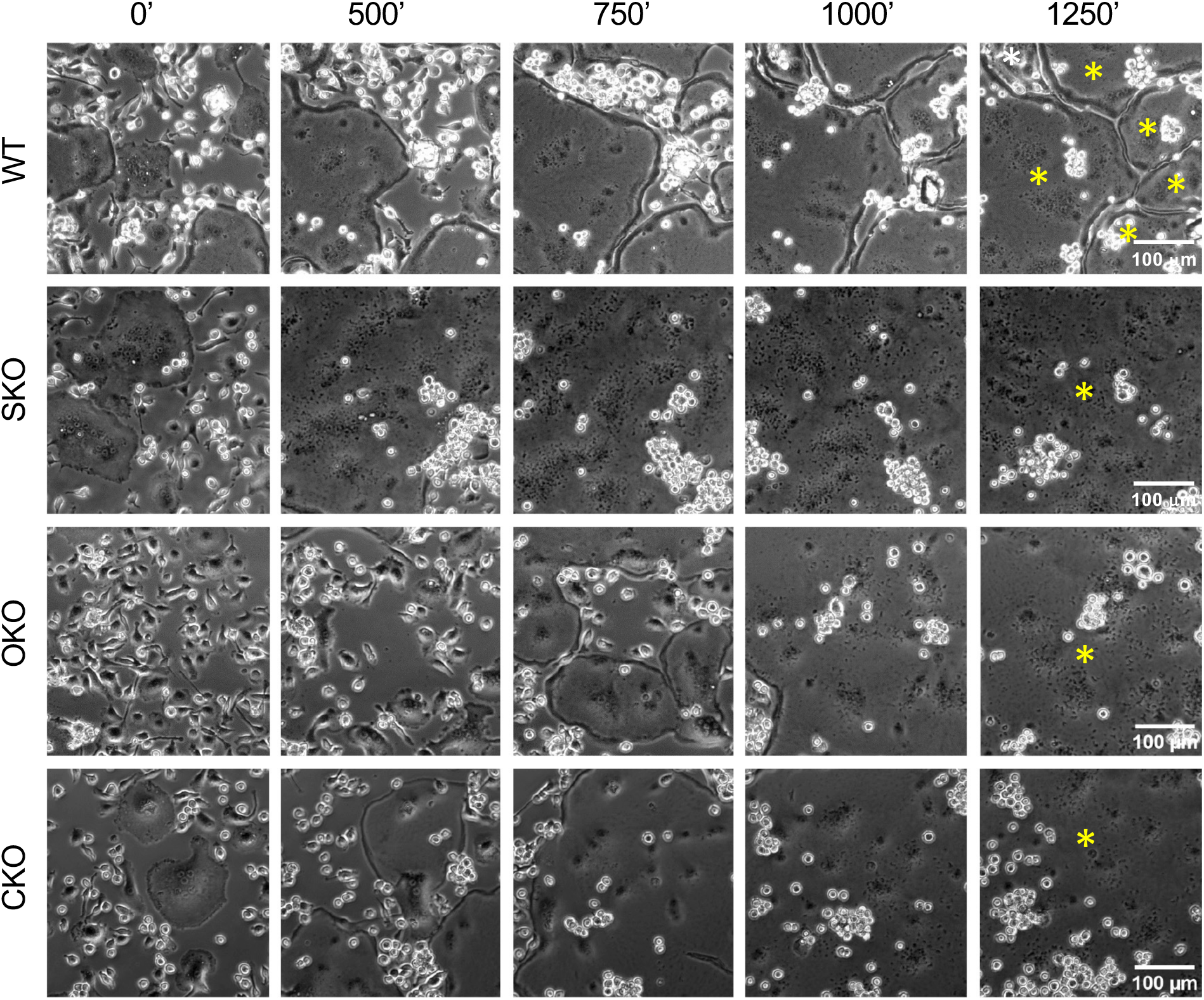
**Image series** showing primary mature WT OCLs juxtaposed for >500 sec without fusing (top), in contrast to mature primary OCLs from SNX10 KO, OSTM1 KO, or CLC-7 KO mice that fuse rapidly. The last frame of the WT series contains 6 OCLs, while the last frame from each KO contains part of a single OCL; each OCL in these frames is marked by a yellow asterisk. Scale bars: 100 μm. Images are taken from movies similar to Movies 1-4.

Differentiation of cells from SKO, OKO, and CKO mice initially proceeded similarly, but mature mutant OCLs continued to fuse with each other, generating progressively larger cells (Fig. 3, Movies 2-4). Fusion persisted until a single large cell occupied the entire growth surface or until no more fusion partners were available, as previously described for SKO OCLs [29].

We next examined the dynamics of fusion between mature OCLs in WT and in the mutant cultures. To this end, we measured the time interval between the initial contact between pairs of cells and their eventual fusion, using phase-contrast video microscopy, as described in Materials and Methods. As shown in Fig. 4 and in Movies 1-4, immature WT OCLs fused with other oligo-nucleated, immature cells 95.3±24.7 minutes (mean±SEM) after making initial contact. In contrast, pairs of mature WT OCLs essentially did not fuse; the cells remained juxtaposed for much longer time periods (1607.0±259.1 minutes; Fig. 4), determined mostly by the time that elapsed from initial contact to separation or to the end of the movie. In contrast, morphologically mature OCLs from the SKO, OKO, and CKO cultures fused readily, with “touch-to-fusion” periods in the range of 75-116 minutes, similar to immature WT cells (Fig. 4). These results indicate that loss of *Snx10*, *Ostm1*, or *Clcn7* drives a largely similar hyper-fusion phenotype, whereby mature OCLs continue to fuse repeatedly with other mature OCLs, forming massive, non-functional cells.

**Figure 4:**
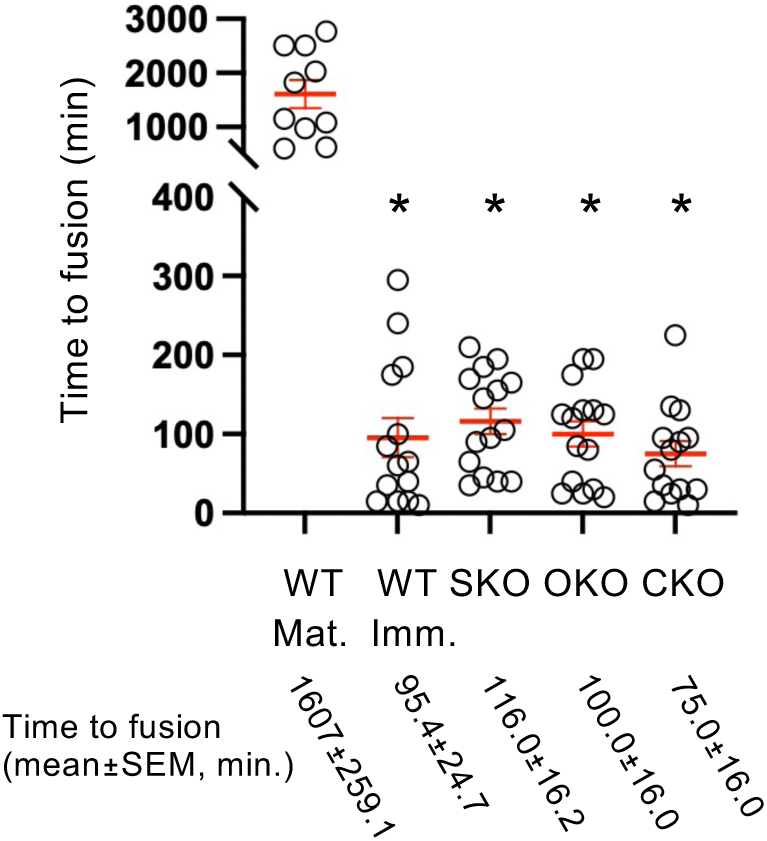
Mature SKO, OKO and CKO OCLs display aberrant fusion kinetics. Shown is the time (mean±SEM) that elapsed from initial contact of two cells to their fusion. WT Mat., WT Imm. – mature and immature WT OCLs, respectively. WT Mature OCLs did not fuse, hence times indicated in that category are from initial contact to end of the movie or to cell separation. *: p≤0.006 relative to WT Mat., by the Kruskal-Wallis multiple comparisons test. WT Imm., SKO, OKO, and CKO groups are not statistically distinct. N=10-15 cell pairs per category.

### Deregulated OCL fusion is not correlated with expression of RANKL-induced genes or with hyper-sensitivity to RANKL

Given RANKL’s crucial role in osteoclastogenesis, we investigated whether deregulated fusion in mutant OCLs stemmed from excessive response to this cytokine. To explore this possibility, we determined the mRNA levels of NFATc1, the major osteoclastogenic transcription factor that is induced by RANKL, and several of its downstream targets in the mutant OCLs relative to WT controls (Fig. 5A). *Nfatc1* levels were reduced in SKO OCLs but unchanged in OKO and CKO OCLs. Expression of *Ctsk*, *Mmp9*, and *Acp5* was largely unaffected, except for slight reductions in OKO and CKO OCLs. Additionally, mRNA levels of *Dcstamp* and *Ocstamp*, which encode major fusion-essential proteins in OCLs, remained unchanged across all mutants (Fig. 5). These results suggest that hyper-fusion in the mutant OCL cultures is not driven by increased responsiveness to RANKL.

**Figure 5:**
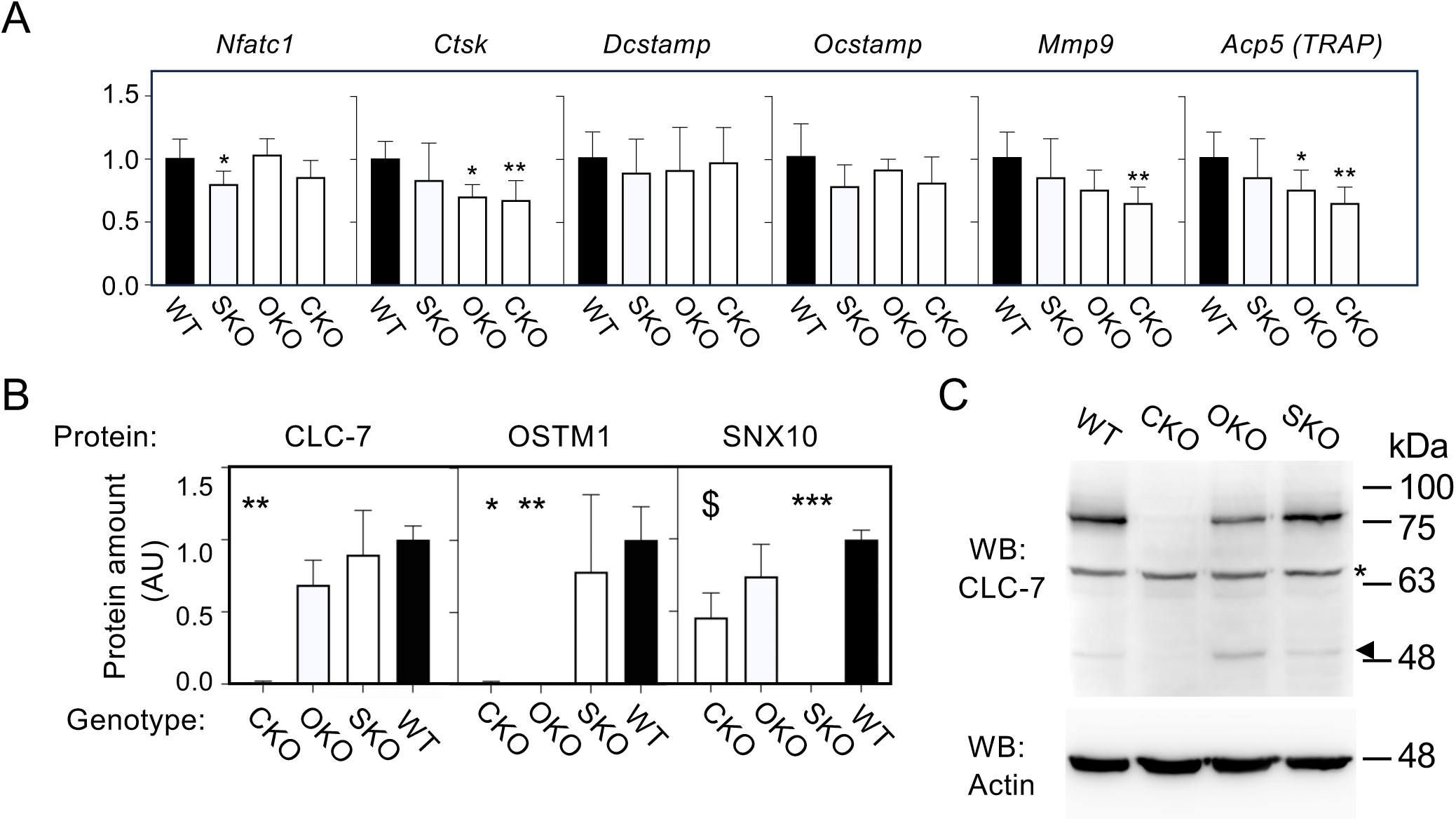
OCL hyperfusion is not caused by excessive RANKL signaling. **A. Quantification by qPCR of the amounts of RANKL-responsive genes** in OCLs cultured from WT, SKO, CKO, and OKO mice. *: p<0.05, **: P<0.01 by the non-parametric Kruskall-Wallis test. N=4-14 independent cultures per bar. **B**. **Relative levels of CLC-7, OSTM1 and SNX10 proteins in OCLs from the strains used in this study.** Protein levels in the various OCL cultures were quantified by targeted mass spectrometry, and are presented relative to levels of the same proteins in WT OCLs (mean±SD). $: p=0.05; *: p=0.019; **: p≤0.0034; ***: p=0.0005, by the non-parametric Kruskall-Wallis test. N=3-6 cultures, each from a separate mouse, per bar. **C**. **Protein blot documenting expression of CLC-7 protein in OCLs of the above genotypes.** Arrowhead denotes probable degradation product of CLC-7. Asterisk denotes non-specific band. WT, wild-type; CKO, CLC-7-KO; OKO, OSTM1-KO; SKO, SNX10-KO.

To further investigate this issue, we examined OCL differentiation at varying RANKL concentrations. Equal numbers of M-CSF-treated splenocytes from WT mice and from the three mutant strains were cultured without RANKL or in the presence of RANKL in concentrations ranging from 2 ng/ml to 200 ng/ml, which represent a ten-fold decrease or increase, respectively, from our standard concentration of 20 ng/ml (Supplementary Fig. 5). No OCLs formed in the absence of RANKL, confirming its necessity for osteoclastogenesis in both WT and mutant cells. OCLs were formed in all the genotypes examined at all concentrations of RANKL, and this occurred more rapidly at higher concentrations of RANKL. Importantly, at all RANKL concentrations, mutant OCLs always exhibited uncontrolled fusion and formed giant cells, whereas WT OCLs underwent regulated fusion, generating smaller OCLs within the normal size range of these cells (Supplementary Fig. 5). These results confirm that while RANKL is essential for fusion, the deregulated nature of fusion in mutant OCLs is independent of the concentration of this cytokine.

CLC-7 protein, but not mRNA, is absent from several tissues of grey-lethal (*gl*) mice, which lack OSTM1, indicating that in some cases OSTM1 is required to stabilize the CLC-7 protein [19]. A trivial explanation for the high phenotypic similarity of OCLs lacking SNX10, OSTM1, or CLC-7 might therefore be that one of these three proteins is missing from all three mutant strains. To examine this possibility, we compared the levels of the SNX10, OSTM1, and CLC-7 proteins in primary OCLs from the three mutants. This study was performed by quantitative mass spectrometry due to the unavailability of antibodies against endogenous SNX10 and OSTM1 that are suitable for protein blotting. As seen in Fig. 5B, knockout of each of the three genes in OCLs resulted in the total absence of its protein product, with small effects on the levels of the other two proteins in most cases. In particular, CLC-7 and OSTM1 proteins were present in WT and in SKO OCLs at similar levels.

Exceptions to this rule occurred in OCLs in which the *Clcn7* gene had been targeted, from which the OSTM1 protein was completely absent, and in which the SNX10 protein was reduced by approximately 50% with borderline (p=0.05) significance (Fig. 5B). This latter result is likely not the cause for the abnormal phenotypes of CKO OCLs, since a similar reduction in SNX10 expression in heterozygous SKO mice does not induce osteopetrosis or abnormal OCL fusion [29].

Examination of CLC-7 protein levels by protein blotting in OCLs from the three mutants confirmed that the CLC-7 protein was prominently present in WT, SKO, and OKO OCLs, and was absent from CKO OCLs (Fig. 5C). In OKO OCLs, the CLC-7 protein band was somewhat degraded (Fig. 5C), in partial agreement with the role of OSTM1 in stabilizing the CLC-7 protein [19]. We conclude that the phenotypic similarities of SKO, OKO, and CKO OCLs are not caused by loss of one of the SNX10, OSTM1, or CLC-7 proteins in all three mutants.

### SNX10 regulates the subcellular trafficking of lysosomes containing OSTM1 and CLC-7 to the cell periphery

Given the striking similarities between the hyper-fusion phenotypes of SKO, OKO, and CKO OCLs, we hypothesized that SNX10, OSTM1, and CLC-7 function within a shared molecular pathway. OSTM1 and CLC-7 form the critical lysosomal Cl⁻/H⁺ antiporter complex [19, 39–42], but their relationship with SNX10 remains unclear. To explore this issue, we determined the respective subcellular distributions of the three proteins, and the molecular interactions between them.

First, we used multi-color immunofluorescence microscopy to visualize the spatial relationships between SNX10, OSTM1, and CLC-7 in WT and mutant OCLs. To this end, RAW 264.7 cells, expressing tagged SNX10, OSTM1, or CLC-7 were prepared. Specifically, an HA tag was inserted into the endogenous *Snx10* gene by CRISPR/CAS9 (Supplementary Fig. 2). A V5-tagged OSTM1 was expressed in OSTM1-knockout cells, and FLAG-tagged CLC-7 was expressed in CLC-7-knockout cells. In the latter two cases, expression of the exogenous tagged proteins restored the regulated fusion phenotype in their respective knockouts, indicating that both tagged proteins retained the function of their WT counterparts (Supplementary Figs. 6, 7). Endogenous CLC-7 was also visualized using specific antibodies against this protein.

We initially compared the pair-wise subcellular localizations of SNX10, OSTM1 or CLC-7 with each other and with the lysosomal protein LAMP1 in WT RAW264.7 OCLs. For further orientation, the cells were also co-stained for actin and DNA, and visualized with 4-color fluorescence microscopy. Examination of the cellular localizations of SNX10 and LAMP1 revealed substantial co-localization of the two proteins in punctate structures located primarily in two distinct subcellular locations, the perinuclear region and the cell periphery (Fig. 6 and Supplementary Fig. 9). The apparent co-localization of SNX10 and LAMP1 in both locations suggested that SNX10 is associated, at least in part, with lysosomes or related LAMP1-containing organelles. Similar perinuclear and cell-peripheral localizations that overlapped with LAMP1 were observed also for OSTM1 and CLC-7 (Fig. 6 and Supplementary Fig. 9), consistent with previous findings [19, 40]. The peripheral arrays of SNX10-, OSTM1- and CLC-7-containing lysosomes were typically flanked by the more peripheral belt of actin-rich of podosomes (Supplementary Fig. 8). Pairwise localization studies confirmed partial colocalization of SNX10 with CLC-7 and with OSTM1 in both the perinuclear region and at the cell periphery (Fig. 7 and Supplementary Figure 10). CLC-7 and OSTM1 displayed strong colocalization at both locations (Fig. 7 and Supplementary Figure 10), in agreement with their previously-documented molecular interactions ([19, 34] and Fig. 9 below).

**Figure 6:**
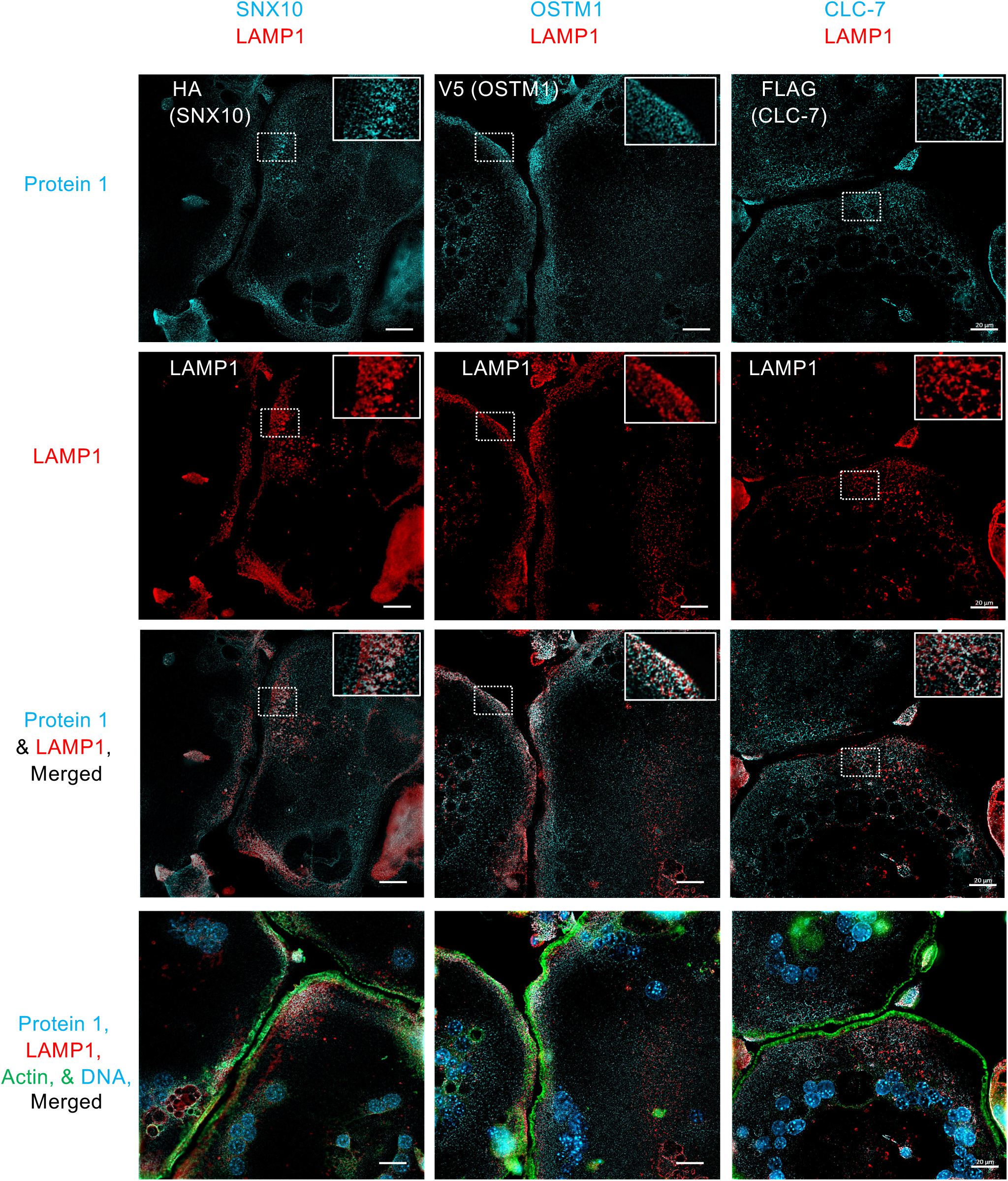
Co-localization of SNX10, OSTM1, or CLC-7 with LAMP1 in OCLs. RAW264.7 cells were differentiated into OCLs and stained as indicated. Dashed rectangles mark areas magnified in the insets at the top right of the figures. As described in the Results, endogenous SNX10 was visualized via a C-terminal HA tag that was inserted into the *Snx10* gene; exogenous, V5-tagged OSTM1 was expressed in OSTM1- KO cells, and exogenous FLAG-tagged CLC-7 was expressed in CLC-7-KO cells. Scale bars: 20 μm.

**Figure 7:**
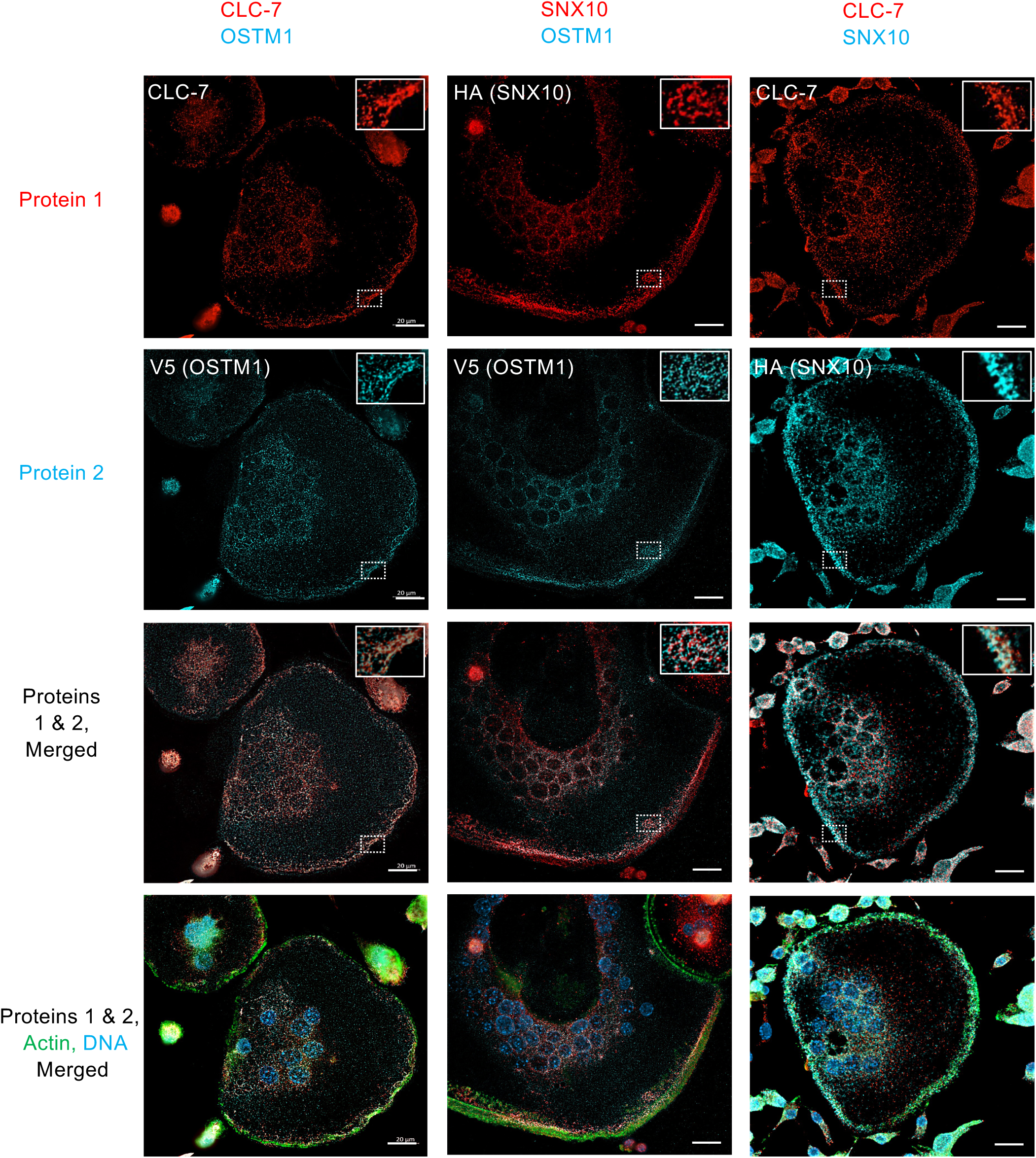
Co-localization of SNX10, OSTM1, and CLC-7 in RAW264.7 OCLs. Dashed rectangles mark areas magnified in the insets at the top right of the figures. Endogenous SNX10 was visualized via a C-terminal HA tag, exogenous OSTM1 was visualized via a C-terminal V5 tag, and endogenous CLC-7 was visualized using anti-CLC-7 antibodies. For SNX10-OSTM1 co-localization, cells expressing SNX10-HA were co-cultured with cells expressing OSTM1-V5; cell fusion produced cells that express both proteins for analysis. Each image represents a single, 0.2 μm-thick Z section. Scale bars: 20 μm. See also Supplementary Figure 8.

Next, we examined the role of SNX10 in the subcellular distribution and colocalization of CLC-7 and OSTM1 by comparing the labeling patterns of these two proteins in WT and SKO OCLs. The presence of both CLC-7 and OSTM1 proteins specifically at the cell periphery was significantly reduced in SKO cells, while the perinuclear expression and localization of these proteins was apparently unaffected (Figs. 8A, 8B). Of note, the overall pattern of LAMP1 staining at the cell periphery was not significantly altered in SKO OCLs (Fig. 8C), suggesting that loss of SNX10 does not affect the presence of other LAMP1-expressing vesicles, such as other lysosomes or late endosomes, at the cell periphery. Collectively, these findings indicate that SNX10 is required for the directed trafficking of CLC-7- and OSTM1-rich vesicles to peripheral regions of mature OCLs.

**Figure 8:**
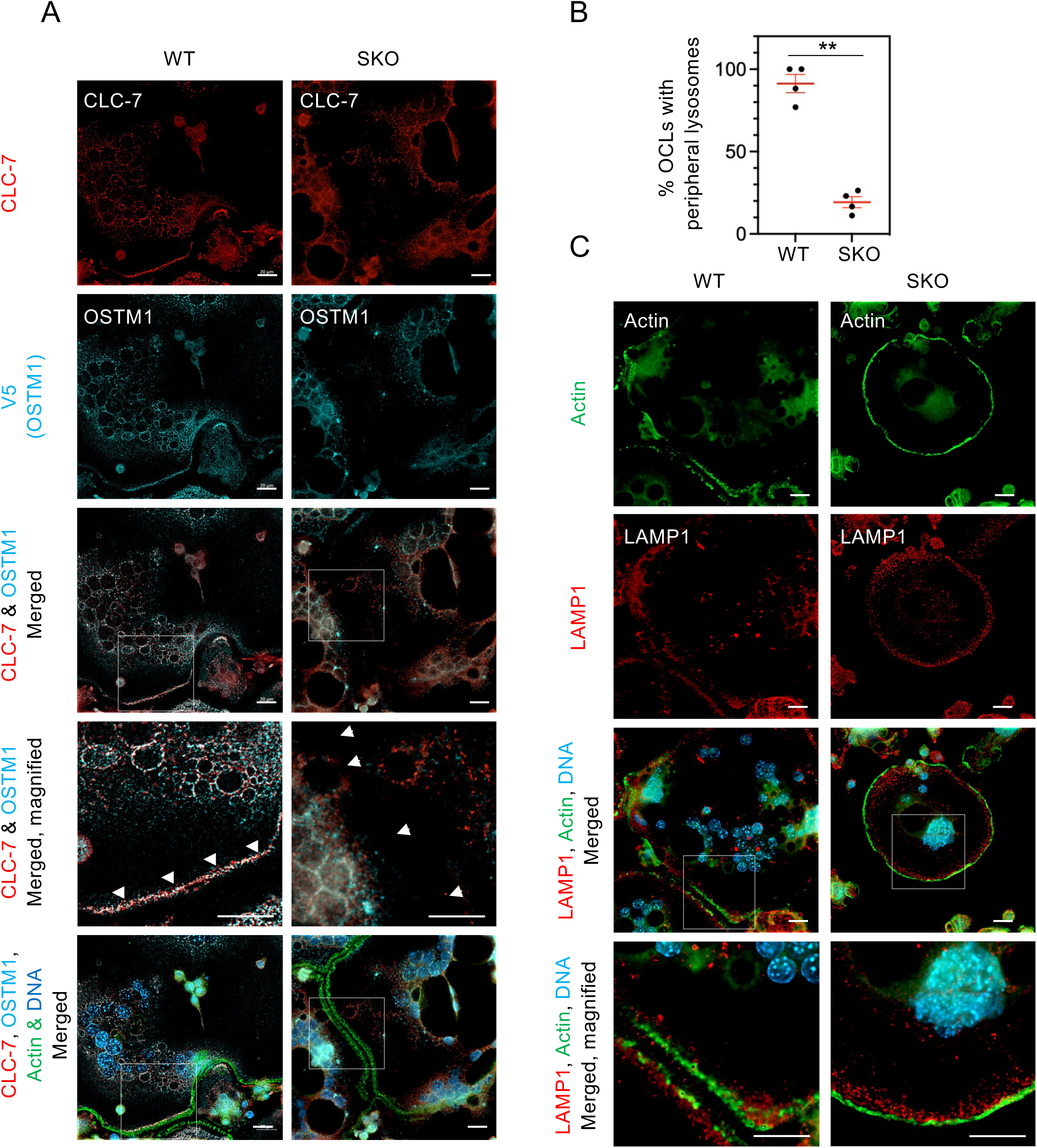
Loss of SNX10 specifically reduces peripheral CLC-7 and OSTM1 in RAW264.7 OCLs. **A. CLC-7 and OSTM1 staining in WT vs. SKO OCLs.** Arrowheads in the merged, magnified images indicate the peripheral CLC-7/OSTM1 (WT cells) or its expected position based on podosome (actin) staining (SKO cells). Both sets of cells express OSTM1-V5, endogenous CLC-7 was visualized using anti-CLC-7 antibodies. **B**. **Percentage of OCLs with peripheral lysosome that contain OSTM1** (mean±SEM). OSTM1-V5 was expressed in WT or SKO OCLs, the cells were stained for V5, and the number of cells with peripheral OSTM1-containing lysosomes was counted. 9-19 cells per experiment and genotype were scored at random from 4 different experiments, for a total of N=50 (WT) or N=53 (SKO) cells. **: p<0.0001 by Students unpaired t test. **C. Peripheral lysosomes are present in SKO OCLs.** Cells were stained for actin and LAMP1. Scale bars: 20 μm.

To explore the molecular mechanisms underlying the differential subcellular distribution of vesicles containing CLC-7 and OSTM1, we next examined whether physical complexes form between SNX10 and CLC-7 or OSTM1. Studies in 293T cells co-expressing these proteins revealed robust co-immunoprecipitation between CLC-7 and SNX10 (Fig. 9A), and similar results were obtained also in OCLs (Fig. 9B). The known interaction between CLC-7 and OSTM1 was observed here as well, and these two proteins co-precipitated in OCL lysates (Fig. 9C). On the other hand, OSTM1 did not co-precipitate with SNX10 (Fig. 9A), suggesting that of these two proteins, CLC-7 is the main interaction partner of SNX10.

**Figure 9:**
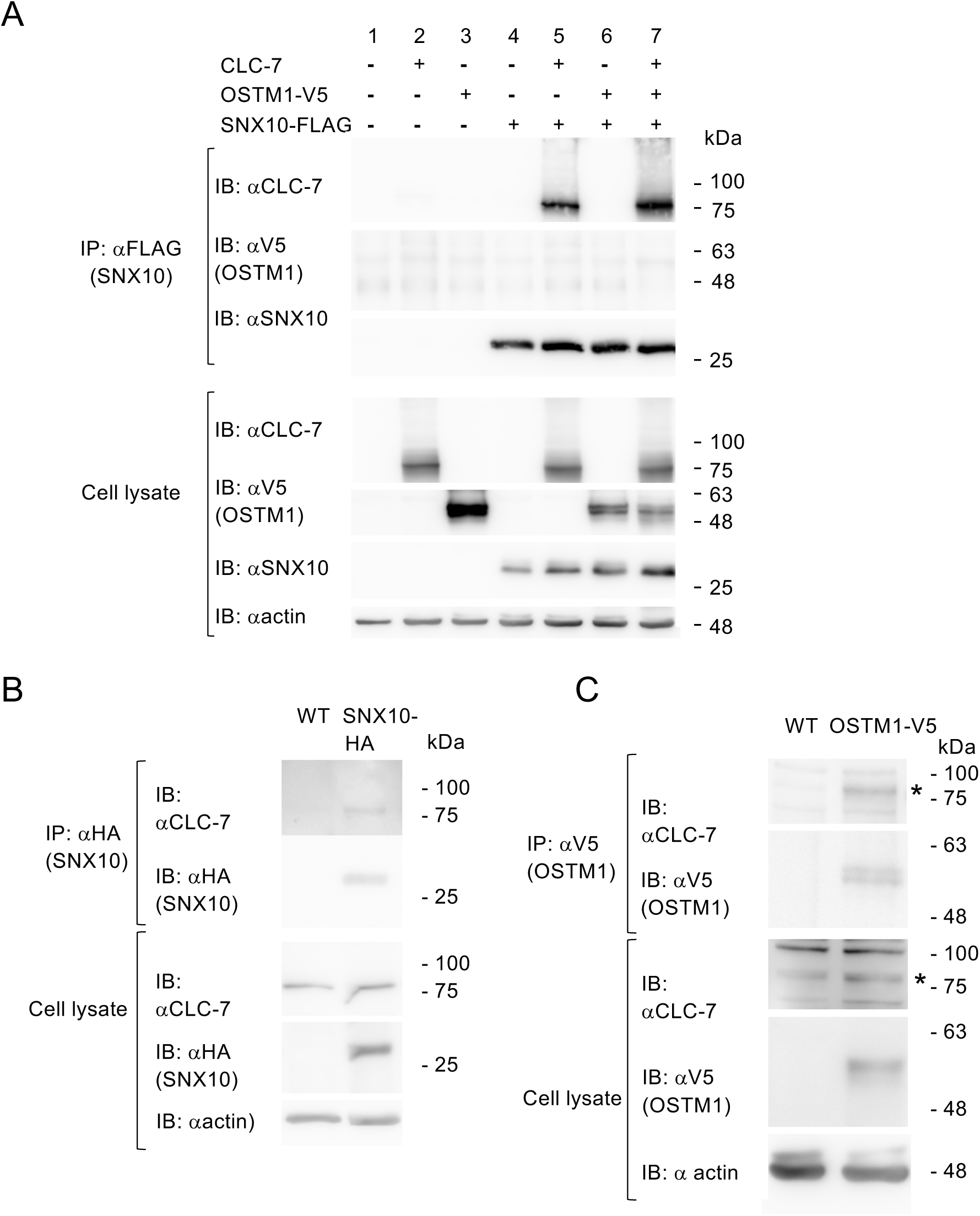
SNX10 co-immunoprecipitates with CLC-7, but not with OSTM1. **A.** Flag-tagged SNX10, V5-tagged OSTM1, and CLC-7 were expressed in HEK293 cells; SNX10 was precipitated via its FLAG tag, and SNX10, OSTM1 and untagged CLC-7 were detected as indicated. **B**. Lysates from wild-type RAW264.7 OCLs, in which endogenous SNX10 was tagged with an HA tag (SNX10-HA) or not (WT), were immunoprecipitated with HA antibodies. Blots were probed with antibodies against HA (SNX10) or CLC-7. **C**. CLC-7 interacts with OSTM1 in RAW264.7 OCLs. Exogenous OSTM1 was expressed (OSTM1-V5) or not (WT) in the cells, and lysates were precipitated with anti-V5 (OSTM1) antibodies. Blots were probed with antibodies against V5 (OSTM1) or CLC-7. Asterisk denotes the CLC-7 protein.

## Discussion

Recent studies of the molecular components associated with ARO provide novel insights into the basic molecular and cellular mechanisms that underlie osteoclast formation and resorptive activity. The key proteins linked to “OCL-rich” ARO fall into two primary categories: those involved in acidification of the resorption lacunae (TCIRG1, CLC-7, OSTM1) and those regulating vesicular trafficking and sorting (primarily SNX10 and PLEKHM1). To characterize the functional interplay between these sorting- and acidification-related proteins, this study focused on SNX10, CLC-7, and OSTM1, addressing two questions: (1) Does the loss of essential components of the “acidification complex” (specifically, CLC-7 and OSTM1) induce deregulated fusion of mature OCLs, similar to the phenotype observed with SNX10 loss? (2) Does SNX10 deficiency affect CLC-7 and OSTM1 in OCLs?

The experiments presented in Figs. 1-4 and in Supplementary Fig. 4 demonstrate that SNX10, CLC-7, and OSTM1 are each essential for both bone resorption by OCLs and for regulating their fusion. These findings suggest that each of these proteins plays critical and non-redundant roles in distinct cellular processes associated with cell fusion and resorption, which likely occur at separate sub-cellular locations - the peripheral plasma membrane and the ruffled border, respectively. Notably, the OCL phenotypes observed in the three mutants were essentially indistinguishable. These include the formation of gigantic OCLs, the deregulated and repeated fusion events between mature OCLs, the similar fusion dynamics (i.e.,, the time between initial contact and actual fusion), the loss of resorption activity, and the formation of a single peripheral podosome belt in mutant OCLs cultured on bone. Interestingly, the fusion dynamics of mature OCLs from all three mutants were not different from those of immature WT OCLs (Fig. 4), suggesting that the nearly complete arrest of fusion of mature WT OCLs is attributable to a robust “fusion stop switch” that is absent from OCLs that lack any of these three proteins. The multiple similarities between the OCL phenotypes of the three mutants strongly suggest that these three proteins function in a concerted manner, most likely within the same cellular pathways. However, the normal levels of OSTM1 and CLC-7 proteins in SKO OCLs (Figs. 5B, 5C) suggests that loss of SNX10 induces the above phenotypes in a manner that is distinct from loss of CLC-7 or OSTM1. A genome-wide genetic screen identified each of the *SNX10*, *OSTM1*, and *CLCN7* genes as essential for the cytotoxic activity of apilimod, an inhibitor of the lipid kinase PIKfyve, in B-cell non-Hodgkin lymphoma [41]. This suggests that these three proteins may function concertedly also in other cellular systems.

The current study highlights the co-localization of SNX10 with CLC-7 and OSTM1 in LAMP1-expressing lysosomes or lysosome-related organelles. Lysosomes are structurally and functionally heterogeneous organelles, whose presence at the cell periphery and in the perinuclear region is well-established [42–45]. Lysosomes present at these two locations differ in their properties (e.g., luminal pH [42]) and in their cellular roles, with perinuclear lysosomes primarily involved in degrading proteins and other cellular components and recycling their building blocks [44, 46]. In contrast, lysosomes located at the cell periphery participate in multiple processes, such as plasma membrane repair [47] and extra-cellular secretion, such as in OCL-driven bone matrix degradation. These latter roles require lysosomes to be present near the plasma membrane and to fuse with it [46]. Trafficking of lysosomes between the perinuclear and cell-peripheral regions is an active and regulated process, which occurs in response to diverse physiological stimuli, such as nutrient starvation [48], oxidative stress [49], or changes in cytosolic pH [50, 51]. Transport of lysosomes in both directions occurs along the microtubular network and is dependent upon different motor proteins: retrograde movement toward the perinuclear region is mediated by dynein/dynactin complexes, while kinesin-based motors facilitate anterograde transport toward the periphery [44, 45].

The co-localization experiments conducted here support the notion that LAMP1- containing lysosomes are heterogeneous in their CLC-7, OSTM1 and SNX10 levels. That said, the co-localization of CLC-7 with OSTM1 in lysosomes is high, in agreement with their known roles in forming the functional H^+^/Cl^-^ exchanger in lysosomes [19, 40]. The finding that CLC-7 protein is expressed in OCLs lacking OSTM1 (Figs. 5B, 5C) is noteworthy in this context. Lange and colleagues showed that the CLC-7 protein was absent from several tissues of grey-lethal (*<u>gl</u>*) mice, which do not express OSTM1, suggesting that OSTM1 is required for stabilization of CLC-7 [19]. Nevertheless, some CLC-7 protein was detected in OCLs from *gl* mice [19]. The finding that the CLC-7 protein is prominently present, yet somewhat degraded, in OKO OCLs (Fig. 5C) agrees with the latter finding, and suggests that, unlike other cell types, presence of CLC-7 protein in OCLs is not entirely dependent on OSTM1. The role of SNX10 in resorption and cell fusion is, however, less clear. Loss of SNX10 did not affect the overall cellular levels of CLC-7 or OSTM1 (Figures 5B, 5C) nor the apparent formation and stabilization of the CLC-7/OSTM1 complex (Figure 8). In contrast, the removal of SNX10 significantly affected the subcellular distribution of CLC-7/OSTM1- containing lysosomes by dramatically reducing their level at the cell periphery. This finding suggests that presence of lysosomes containing CLC-7 and OSTM1 in this region is essential for the physiological regulation of both cell fusion and bone resorption. Loss of CLC-7 or OSTM1, or mis-localization of lysosomes containing these proteins through loss of SNX10, can lead to similar functional outcomes.

Co-immunoprecipitation data presented here indicates that CLC-7 is an interaction partner of SNX10, while OSTM1 is not (Fig. 9). OSTM1 may associate with SNX10 indirectly through its association with CLC-7. Yet, OSTM1 did not precipitate with SNX10 even when CLC-7 was present, possibly due to the added difficulty in demonstrating indirect associations. Alternatively, association of CLC-7 with SNX10 and with OSTM1 might be mutually-exclusive events. This would define two sub-populations of CLC-7, with the CLC- 7-SNX10 sub-population possibly involved in regulating lysosome transport. Further studies are required to clarify this point. Prior studies have shown that OSTM1 physically associates in osteoclasts with KIF5B [34], a member of the kinesin 1 family that participates in the trafficking of lysosomes to the cell periphery [44]. Knockdown of KIF5B in COS7 cells enhanced perinuclear clustering of lysosomes and of OSTM1 and CLC-7 proteins [34], while loss of OSTM1 in OCLs disrupted the localization, but not acidification, of lysosomes [25]. These finding suggest that the OSTM1-KIF5B interaction is essential for the proper cellular distribution of lysosomes and of OSTM1/CLC-7 within the lysosomal system. Along these lines, we propose that SNX10 stabilizes interactions between lysosomal proteins and the motor protein networks that drive anterograde transport along the microtubular network. Given its PX domain, which binds PI3P and PI(3,5)P2 [33], SNX10 may anchor to the lysosomal membrane and facilitate interactions with CLC-7 and other proteins. KIF5B does not bind CLC-7 [34] and SNX10 does not bind OSTM1 (Fig. 9), suggesting that CLC-7 and OSTM1 interact with distinct proteins in this context.

Transport of lysosomes towards the plasma membrane and their eventual fusion with it are essential for the ability of OCLs to secrete protons and proteases during bone degradation. Loss of peripheral lysosomes is then consistent with the inactivity of the mutant OCLs. A question that remains open is whether loss of peripheral lysosomes that contain CLC-7 and OSTM1 also affects the regulation of OCL fusion. Lysosomes contribute to membrane protein removal, degradation, and recycling, so their disruption may interfere with these processes, potentially trapping fusion-promoting proteins at the cell surface. Supporting this, OCLs homozygous for the ARO-inducing R51Q SNX10 mutation, which lack SNX10 protein and exhibit continuous fusion similar to SKO OCLs, display reduced endocytosis and increased levels of DC-STAMP at the cell periphery [33, 29]. Alternatively, loss of peripheral lysosomes may impair plasma membrane repair, possibly promoting contact and fusion between adjacent membranes. Further studies are required to explore these mechanisms.

When cultured on bone, the giant OCLs lacking SNX10, OSTM1 or CLC-7 each display a single peripheral ring of podosomes (Fig. 2), a characteristic feature of OCLs that are in contact with non-resorbable surfaces such as plastic or glass. This finding strongly suggests that the mutant OCLs fail to recognize the underlying bone surface as such, potentially indicating a defect in integrin-mediated bone recognition. Further studies are required to directly examine this issue and to determine whether it is linked with the hyper-fusion phenotype of these cells.

Collectively, our findings suggest that SNX10, OSTM1, and CLC-7 function together to generate active OCLs of appropriate functional sizes. We propose that OSTM1 and CLC-7 are essential for these processes, and that loss of either protein or their mis-localization through loss of SNX10 similarly disrupt the osteoclastogenic process and generate the same complex phenotype of inactive, continuously-fusing OCLs.

## Acknowledgements

We thank Prof. Thomas Jentsch, Leibniz-Forschungsinstitut für Molekulare Pharmakologie, Berlin, Germany, for kind gifts of CLC-7 knockout mice and anti-CLC-7 antibodies. We also thank Ms. Ofira Higfa and Mr. Neriah Sharabi of the Weizmann Institute Department of Veterinary Services for expert animal care. This study was supported by the Israel Science Foundation (grant #1734/20, to AE), by the Kekst Family Institute for Medical Genetics of the Weizmann Institute of Science (to AE) and by the Canadian Institute of Health Research (CIHR), grant #86655 (to JV). AE is the incumbent of the Marshall and Renette Ezralow Professorial Chair.

## Movie legends

**Movie 1:** Phase contrast light microscopy-based live cell imaging of WT monocytes grown on optical grade plastic as they fuse into OCLs. Individual images were obtained at 5-minute intervals for a period of 22 hours. Note the formation of large mature OCLs that do not further fuse with each other. Still images from this movie are presented in Figure 3.

**Movie 2:** Phase contrast light microscopy-based live cell imaging of SKO (SNX10-KO) monocytes grown on optical grade plastic as they fuse into OCLs. Individual images were obtained at 5-minute intervals for a period of 28.5 hours. Note extensive and continuous fusion between mature OCLs that results in formation of a large cell that eventually covers the entire field of view. Still images from this movie are presented in Figure 3.

**Movie 3:** Phase contrast light microscopy-based live cell imaging of OKO (OSTM1-KO) monocytes grown on optical grade plastic as they fuse into OCLs. Individual images were obtained at 5-minute intervals for a period of 22.5 hours. Note extensive and continuous fusion between mature OCLs that results in formation of a large cell that eventually covers most of the field of view. Still images from this movie are presented in Figure 3.

**Movie 4:** Phase contrast light microscopy-based live cell imaging of CKO (CLC-7-KO) monocytes grown on optical grade plastic as they fuse into OCLs. Individual images were obtained at 5-minute intervals for a period of 20 hours. Note extensive and continuous fusion between mature OCLs that results in formation of a large cell that eventually covers the entire field of view. Still images from this movie are presented in Figure 3.

## References

1. Charles JF, Aliprantis AO. Osteoclasts: more than ‘bone eaters’. Trends Mol Med. 2014;20(8):449–459. doi: 10.1016/j.molmed.2014.06.001.

2. Feng X, Teitelbaum SL. Osteoclasts: New Insights. Bone Res. 2013;1(1):11–26. doi: 10.4248/br201301003.

3. Teitelbaum SL. Osteoclasts: what do they do and how do they do it? Am J Pathol. 2007;170(2):427–435. doi: 10.2353/ajpath.2007.060834

4. Elson A, Anuj A, Barnea-Zohar M, Reuven N. The origins and formation of bone-resorbing osteoclasts. Bone. 2022;164:116538. doi: 10.1016/j.bone.2022.116538.

5. Tsukasaki M, Huynh NC, Okamoto K, Muro R, Terashima A, Kurikawa Y, et al. Stepwise cell fate decision pathways during osteoclastogenesis at single-cell resolution. Nat Metab. 2020;2(12):1382–1390. doi: 10.1038/s42255-020-00318-y.

6. Teitelbaum SL. The osteoclast and its unique cytoskeleton. Ann N Y Acad Sci. 2011;1240:14–17. doi: 10.1111/j.1749-6632.2011.06283.x.

7. Novack DV, Teitelbaum SL. The osteoclast: friend or foe? Annu Rev Pathol. 2008;3:457–484. doi: 10.1146/annurev.pathmechdis.3.121806.151431

8. Akhtari M, Mansuri J, Newman KA, Guise TM, Seth P. Biology of breast cancer bone metastasis. Cancer Biol Ther. 2008;7(1):3–9. doi: 10.4161/cbt.7.1.5163.

9. Palagano E, Menale C, Sobacchi C, Villa A. Genetics of Osteopetrosis. Curr Osteoporos Rep. 2018;16(1):13–25. doi: 10.1007/s11914-018-0415-2.

10. Sobacchi C, Schulz A, Coxon FP, Villa A, Helfrich MH. Osteopetrosis: genetics, treatment and new insights into osteoclast function. Nat Rev Endocrinol. 2013;9(9):522–536. doi: 10.1038/nrendo.2013.137.

11. Polgreen LE, Imel EA, Econs MJ. Autosomal dominant osteopetrosis. Bone. 2023;170:116723.

12. Penna S, Villa A, Capo V. Autosomal recessive osteopetrosis: mechanisms and treatments. Dis Model Mech. 2021;14(5).

13. Kornak U, Schulz A, Friedrich W, Uhlhaas S, Kremens B, Voit T, et al. Mutations in the a3 subunit of the vacuolar H(+)-ATPase cause infantile malignant osteopetrosis. Hum Mol Genet. 2000;9(13):2059–2063. doi: 10.1093/hmg/9.13.2059.

14. Frattini A, Orchard PJ, Sobacchi C, Giliani S, Abinun M, Mattsson JP, et al. Defects in TCIRG1 subunit of the vacuolar proton pump are responsible for a subset of human autosomal recessive osteopetrosis. Nat Genet. 2000;25(3):343–346. doi: 10.1038/77131.

15. Frattini A, Pangrazio A, Susani L, Sobacchi C, Mirolo M, Abinun M, et al. Chloride channel ClCN7 mutations are responsible for severe recessive, dominant, and intermediate osteopetrosis. J Bone Miner Res. 2003;18(10):1740–1747. doi: 10.1359/jbmr.2003.18.10.1740.

16. Quarello P, Forni M, Barberis L, Defilippi C, Campagnoli MF, Silvestro L, et al. Severe malignant osteopetrosis caused by a GL gene mutation. J Bone Miner Res. 2004;19(7):1194–1199. doi: 10.1359/jbmr.040407.

17. Ramírez A, Faupel J, Goebel I, Stiller A, Beyer S, Stöckle C, et al. Identification of a novel mutation in the coding region of the grey-lethal gene OSTM1 in human malignant infantile osteopetrosis. Hum Mutat. 2004;23(5):471–476. doi: 10.1002/humu.20028.

18. Kornak U, Kasper D, Bosl MR, Kaiser E, Schweizer M, Schulz A, et al. Loss of the ClC- 7 chloride channel leads to osteopetrosis in mice and man. Cell. 2001;104(2):205–215. doi: 10.1016/s0092-8674(01)00206-9

19. Lange PF, Wartosch L, Jentsch TJ, Fuhrmann JC. ClC-7 requires Ostm1 as a beta-subunit to support bone resorption and lysosomal function. Nature. 2006;440(7081):220–223. doi: 10.1038/nature04535.

20. Pangrazio A, Poliani PL, Megarbane A, Lefranc G, Lanino E, Di Rocco M, et al. Mutations in OSTM1 (grey lethal) define a particularly severe form of autosomal recessive osteopetrosis with neural involvement. J Bone Miner Res. 2006;21(7):1098–1105. doi: 10.1359/jbmr.060403.

21. Chalhoub N, Benachenhou N, Rajapurohitam V, Pata M, Ferron M, Frattini A, et al. Grey-lethal mutation induces severe malignant autosomal recessive osteopetrosis in mouse and human. Nat Med. 2003;9(4):399–406. doi: 10.1038/nm842.

22. Aker M, Rouvinski A, Hashavia S, Ta-Shma A, Shaag A, Zenvirt S, et al. An SNX10 mutation causes malignant osteopetrosis of infancy. J Med Genet. 2012;49(4):221–226. doi: 10.1136/jmedgenet-2011-100520.

23. Neutzsky-Wulff AV, Sims NA, Supanchart C, Kornak U, Felsenberg D, Poulton IJ, et al. Severe developmental bone phenotype in ClC-7 deficient mice. Dev Biol. 2010;344(2):1001–1010. doi: 10.1016/j.ydbio.2010.06.018.

24. Weinert S, Jabs S, Supanchart C, Schweizer M, Gimber N, Richter M, et al. Lysosomal pathology and osteopetrosis upon loss of H+-driven lysosomal Cl- accumulation. Science. 2010;328(5984):1401–1403. doi: 10.1126/science.1188072.

25. Pata M, Vacher J. Ostm1 Bifunctional Roles in Osteoclast Maturation: Insights From a Mouse Model Mimicking a Human OSTM1 Mutation. J Bone Miner Res. 2018;33(5):888–898. doi: 10.1002/jbmr.3378.

26. Rajapurohitam V, Chalhoub N, Benachenhou N, Neff L, Baron R, Vacher J. The mouse osteopetrotic grey-lethal mutation induces a defect in osteoclast maturation/function. Bone. 2001;28(5):513–523. doi: 10.1016/s8756-3282(01)00416-1.

27. Stein M, Barnea-Zohar M, Shalev M, Arman E, Brenner O, Winograd-Katz S, et al. Massive osteopetrosis caused by non-functional osteoclasts in R51Q SNX10 mutant mice. Bone. 2020;136:115360. doi: 10.1016/j.bone.2020.115360.

28. Ye L, Morse LR, Zhang L, Sasaki H, Mills JC, Odgren PR, et al. Osteopetrorickets due to Snx10 deficiency in mice results from both failed osteoclast activity and loss of gastric acid-dependent calcium absorption. PLoS Genet. 2015;11(3):e1005057. doi: 10.1371/journal.pgen.1005057.

29. Barnea-Zohar M, Stein M, Reuven N, Winograd-Katz S, Lee S, Addadi Y, et al. SNX10 regulates osteoclastogenic cell fusion and osteoclast size in mice. J Bone Miner Res. 2024. doi: 10.1093/jbmr/zjae125.

30. Kasper D, Planells-Cases R, Fuhrmann JC, Scheel O, Zeitz O, Ruether K, et al. Loss of the chloride channel ClC-7 leads to lysosomal storage disease and neurodegeneration. Embo j. 2005;24(5):1079–1091. doi: 10.1038/sj.emboj.7600576.

31. Héraud C, Griffiths A, Pandruvada SN, Kilimann MW, Pata M, Vacher J. Severe neurodegeneration with impaired autophagy mechanism triggered by ostm1 deficiency. J Biol Chem. 2014;289(20):13912–13925. doi: 10.1074/jbc.M113.537233.

32. Vacher J. OSTM1 pleiotropic roles from osteopetrosis to neurodegeneration. Bone. 2022;163:116505. doi: 10.1016/j.bone.2022.116505.

33. Barnea-Zohar M, Winograd-Katz SE, Shalev M, Arman E, Reuven N, Roth L, et al. An SNX10-dependent mechanism downregulates fusion between mature osteoclasts. J Cell Sci. 2021;134(9). doi: 10.1242/jcs.254979.

34. Pandruvada SN, Beauregard J, Benjannet S, Pata M, Lazure C, Seidah NG, et al. Role of Ostm1 Cytosolic Complex with Kinesin 5B in Intracellular Dispersion and Trafficking. Mol Cell Biol. 2016;36(3):507–521. doi: 10.1128/mcb.00656-15.

35. Anuj A, Reuven N, Roberts SGE, Elson A. BASP1 down-regulates RANKL-induced osteoclastogenesis. Exp Cell Res. 2023;431(1):113758. doi: 10.1016/j.yexcr.2023.113758.

36. Schindelin J, Arganda-Carreras I, Frise E, Kaynig V, Longair M, Pietzsch T, et al. Fiji: an open-source platform for biological-image analysis. Nat Methods. 2012;9(7):676–682. doi: 10.1038/nmeth.2019.

37. Livak KJ, Schmittgen TD. Analysis of relative gene expression data using real-time quantitative PCR and the 2(-Delta Delta C(T)) Method. Methods. 2001;25(4):402–408. doi: 10.1006/meth.2001.1262.

38. Gil-Henn H, Elson A. Tyrosine phosphatase-epsilon activates Src and supports the transformed phenotype of Neu-induced mammary tumor cells. J Biol Chem. 2003;278(18):15579–15586.

39. Varnavides G, Madern M, Anrather D, Hartl N, Reiter W, Hartl M. In Search of a Universal Method: A Comparative Survey of Bottom-Up Proteomics Sample Preparation Methods. J Proteome Res. 2022;21(10):2397–2411.

40. Leisle L, Ludwig CF, Wagner FA, Jentsch TJ, Stauber T. ClC-7 is a slowly voltage-gated 2Cl(-)/1H(+)-exchanger and requires Ostm1 for transport activity. EMBO J. 2011;30(11):2140–2152.

41. Gayle S, Landrette S, Beeharry N, Conrad C, Hernandez M, Beckett P, et al. Identification of apilimod as a first-in-class PIKfyve kinase inhibitor for treatment of B-cell non-Hodgkin lymphoma. Blood. 2017;129(13):1768–1778. doi: 10.1182/blood-2016-09-736892.

42. Johnson DE, Ostrowski P, Jaumouillé V, Grinstein S. The position of lysosomes within the cell determines their luminal pH. J Cell Biol. 2016;212(6):677–692.

43. Saffi GT, Botelho RJ. Lysosome Fission: Planning for an Exit. Trends Cell Biol. 2019;29(8):635–646. doi: 10.1016/j.tcb.2019.05.003.

44. Cabukusta B, Neefjes J. Mechanisms of lysosomal positioning and movement. Traffic. 2018;19(10):761–769.

45. Bonifacino JS, Neefjes J. Moving and positioning the endolysosomal system. Curr Opin Cell Biol. 2017;47:1–8.

46. Bouhamdani N, Comeau D, Turcotte S. A Compendium of Information on the Lysosome. Front Cell Dev Biol. 2021;9:798262.

47. Corrotte M, Castro-Gomes T. Lysosomes and plasma membrane repair. Curr Top Membr. 2019;84:1–16. doi: 10.1016/bs.ctm.2019.08.001.

48. Korolchuk VI, Saiki S, Lichtenberg M, Siddiqi FH, Roberts EA, Imarisio S, et al. Lysosomal positioning coordinates cellular nutrient responses. Nat Cell Biol. 2011;13(4):453–460.

49. Sasazawa Y, Souma S, Furuya N, Miura Y, Kazuno S, Kakuta S, et al. Oxidative stress-induced phosphorylation of JIP4 regulates lysosomal positioning in coordination with TRPML1 and ALG2. EMBO J. 2022;41(22):e111476.

50. Heuser J. Changes in lysosome shape and distribution correlated with changes in cytoplasmic pH. J Cell Biol. 1989;108(3):855–864.

51. Parton RG, Dotti CG, Bacallao R, Kurtz I, Simons K, Prydz K. pH-induced microtubule-dependent redistribution of late endosomes in neuronal and epithelial cells. J Cell Biol. 1991;113(2):261–274.

