## Supplemental for "The molecular and cellular interplay between the osteopetrosis-associated proteins SNX10, OSTM1, and CLC-7 during osteoclastogenesis"

Supplemental Tables and Figures

| **Gene** | **Forward** | **Reverse** | **Region** |
|---|---|---|---|
| *Clcn7* for KO | TCCCTCCTTTTCTCCCCCAA | GGAGCACGTATCTGCCCTAC | Exon 5 |
| *Ostm1* for KO | CCATTGAGCTGTCAACCGGA | GTCCATTTCGCTCCGACAGA | Exon 5 |
| *Snx10* for KO | GCTCGTGTGTGTTTCTCACG | AACACTTCTGGGGGCCATTC | Exon 4 |
| *Snx10* for KI (HA tag) | CATCGGGGCGAAGTTCCCTA Primer upstream of homology arm | GCCCTATCTGGCTTTAAATGCTACT | Exon 7 |

Supplemental Table 1: Primers used to amplify genomic regions targeted by CRISPR guides.

| Gene | Forward | Reverse |
|---|---|---|
| *Acp5*(TRAP) | GATGCCAGCGACAAGAGGTT | CATACCAGGGGATGTTGCGAA |
| *Clcn7* set 1 | GACTGGCTGTGGGAAAGGAA | TCTCGCTTGAGTGATGTTGACC |
| *Clcn7* set g* | TGGAGATCAAGCGCTGGGTT | CCGCCCTTCTCTGTGAACTT |
| *Ctsk* | GGCCAGGATGAAAGTTGTATGTATAA | TCTCGTTCCCCACAGGAATC |
| *Dc-stamp* | TCCTCCATGAACAAACAGTTCCA | AGACGTGGTTTAGGAATGCAGCTC |
| *Mmp9* | CTGGACAGCCAGACACTAAAG | CTCGCGGCAAGTCTTCAGAG |
| *Nfatc1* | CCTGGAGATCCCGTTGCTT | CGATGTCTGTCTCCCCTTTCC |
| *Oc-stamp* | TCACAGTCAAATATGACGCCTCAT | GGTGGTTGAGCCTGTGGTAGA |
| *Ostm1* | GTGGTTGCTGTGTCTGTGTTCA | ACGTTTGGGTAGAATGAGTTTGC |
| *Rpl4* | GATGAGCTGTATGGCACTTGG | CTTGTGCATGGGCAGGTTA |
| *Snx10* | ACGCGTTGCTGGTACAATTACC | AAGAGGTGGAGGCTGCTATCG |

* The forward primer overlaps the deletion in the CRISPR *Clcn7* KO cells. Therefore, mutated *Clcn7* transcripts are not detected by this primer set.

Supplemental Table 2: Primers used in qPCR experiments in this study.

| Plain Sequence | Gene | Protein | UniprotID |
|---|---|---|---|
| IGQMNNVELDDELLDPEVDPPHTFPK | Clcn7 | H(+)/Cl(-) exchange transporter 7 (CLC-7) | O70496 |
| YESLDYDNSENQLFLEEER | Clcn7 | H(+)/Cl(-) exchange transporter 7 (CLC-7) | O70496 |
| EVMSTPVTCLR | Clcn7 | H(+)/Cl(-) exchange transporter 7 (CLC-7) | O70496 |
| LQGLILR | Clcn7 | H(+)/Cl(-) exchange transporter 7 (CLC-7) | O70496 |
| FPPIQSIHVSQDER | Clcn7 | H(+)/Cl(-) exchange transporter 7 (CLC-7) | O70496 |
| LQSNALLVQLPELPSK | Snx10 | Sorting nexin-10 | Q9CWT3 |
| QGLEDFLR | Snx10 | Sorting nexin-10 | Q9CWT3 |
| YSVEEAIHK | Snx10 | Sorting nexin-10 | Q9CWT3 |
| NLFFNMNNR | Snx10 | Sorting nexin-10 | Q9CWT3 |
| LCQTCYPLFQQVAIK | Ostm1 | Osteopetrosis-associated transmembrane protein 1 (OSTM1) | Q8BGT0 |
| SSTSFANIQENAT | Ostm1 | Osteopetrosis-associated transmembrane protein 1 (OSTM1) | Q8BGT0 |

Supplemental Table 3: Peptides targeted for detection of CLC-7, OSTM1, and SNX10 in MS studies.

A

| Gene | Chr # | Protein | Targeted exon | Guide sequence | | Outcome | # Independent clones isolated |
|---|---|---|---|---|---|---|---|
| ***Snx10*** | 6 | SNX10 | 4 | AAACATCTTGTGTACGAAGA | AGG | Indel, loss of reading frame (both alleles) | 12 |
| ***Ostm1*** | 10 | OSTM1 | 5 | TTTACAGTTTCTGCACACTT | CGG | Indel, loss of reading frame (both alleles) | 5 |
| ***Clcn7*** | 17 | CLCN7 | 5 | GGACAGTGGAGATCAAGCGC | TGG | Indel, loss of reading frame (both alleles) | 4 |

B

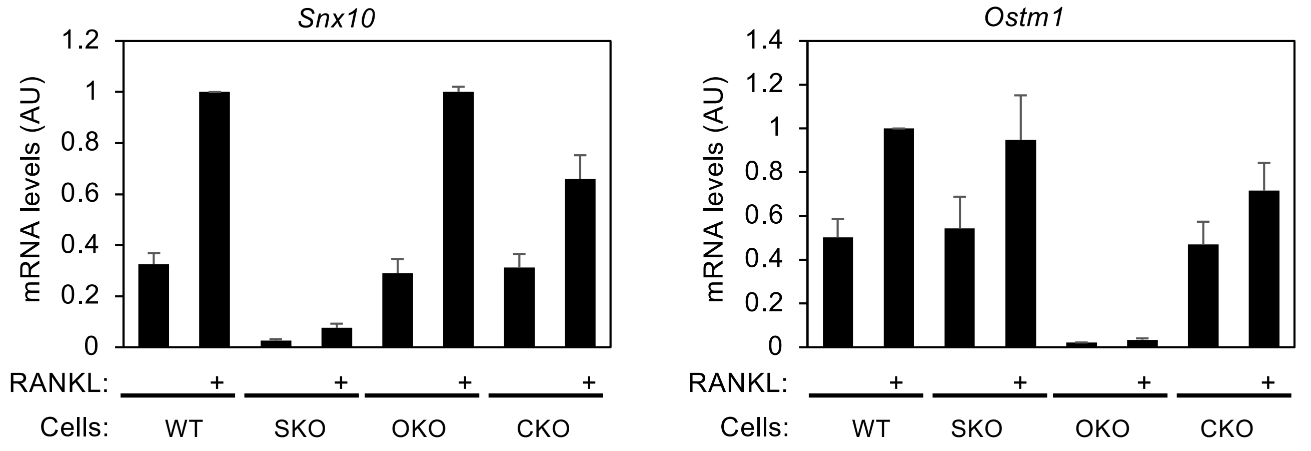


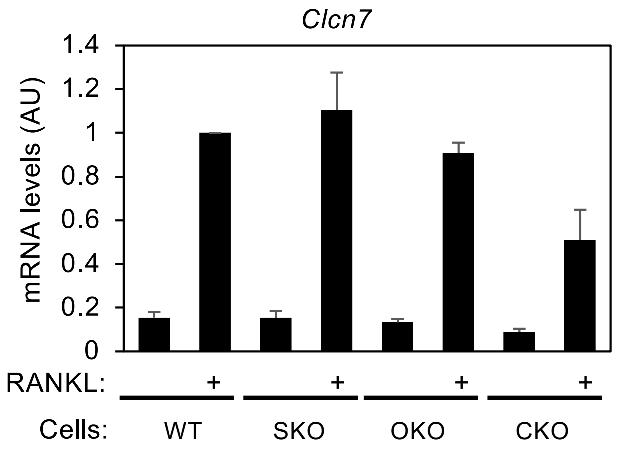


C

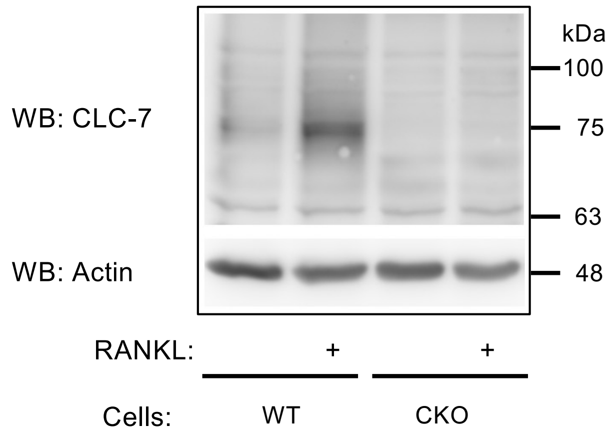


**Supplementary Figure 1: Knocking out *Snx10, Ostm1*, and *Clcn7* in RAW264.7 cells.**

**A.** Table depicting the details of the production of RNA 267.4 clones in which *Snx10*, *Ostm1*, or *Clcn7* (SKO, OKO, CKO, respectively) had been targeted by CRISPR. In most cases each clone contained two distinct loss of reading frame alleles. Production of SKO clones was described in (28) and is shown for completeness of presentation. **B.** mRNA levels of *Snx10, Ostm1*, and *Clcn7* were determined by qPCR in representative clones of RAW264.7 cells in which the *Snx10* (clone La), *Ostm1* (clone L1) or *Clcn7* (clone L11A) genes had been targeted. Data are normalized to mRNA levels of WT, RANKL-treated cells. Data are mean±SD, N=3 independent experiments per bar. **C**. Protein blot of *Clcn7* KO clone L11A, showing the absence of CLC-7 protein despite the presence of *Clcn7* mRNA.

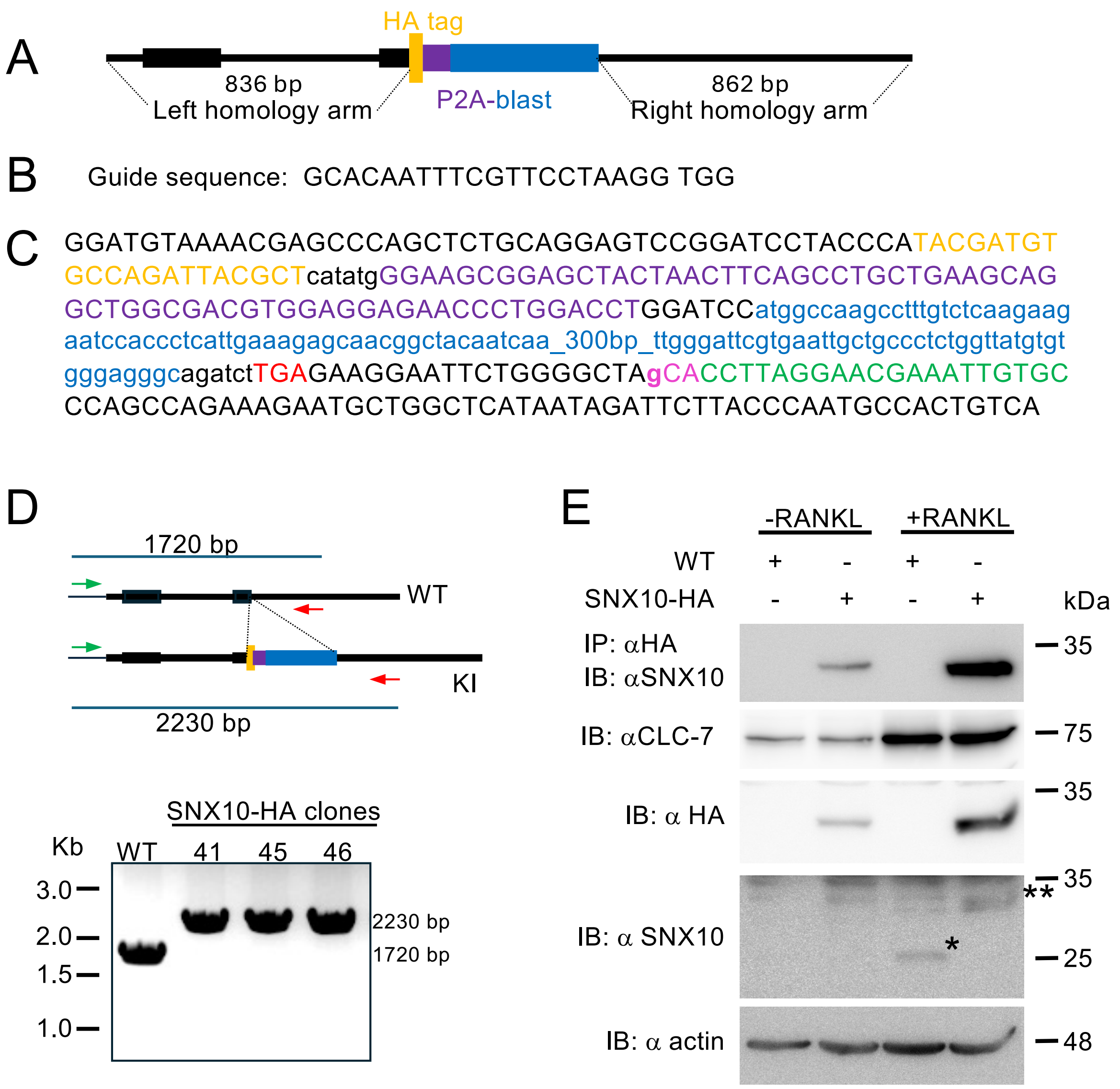


**Supplementary Figure 2. CRISPR-mediated tagging of endogenous SNX10**. A) Schematic representation of the donor DNA construct that was cloned into the pBluescript backbone. The last coding exons of SNX10 are represented by black rectangles, while introns and non-coding sequences are thinner black lines.The HA tag (yellow) follows the last coding exon of SNX10, separated by a 4aa linker. The HA tag is followed by a P2A sequence (purple), the coding region of a blasticidin resistance gene (blue), and the SNX10 termination codon (red in Panel C). The P2A peptide ensures that SNX10-HA and blasticidin resistance will be expressed from the same promoter, but as separate proteins. B) CRISPR guide used to target *Snx10* downstream of the HA-P2A-blast insertion site. C) The sequence of the insert, flanked by the SNX10 endogenous DNA. Sequences are colored as in Panel A. The sgRNA sequence is marked in green (note that the sgRNA is the complementary strand). The PAM sequence (pink) was mutated in the donor DNA (lowerase g from the original C) to eliminate the NGG PAM sequence. D) TOP: Scheme showing PCR amplification strategy of Genomic DNA from WT and SNX10-HA knock-in (KI) clones using a forward primer upstream of the left homology arm (green), and a reverse primer downstream of the insert (red). Primer sequences are shown in Supplemental Table 1. Bottom: Amplification products from three different knock-in clones are shown: 41, 45, and 46. The knock-in clones exhibited only the 2230bp band, and no other bands, indicating either biallelic insertion or a large deletion in the second allele. Sequencing of the knock-in PCR products validated the clones (not shown). E) WT and SNX10-HA clone 45 cells were grown without or with RANKL. Cell extracts were subjected to immunoprecipitation with anti-HA agarose, and IP and total cell lysates were analyzed by SDS-PAGE and protein blotting with the indicated antibodies. The anti-SNX10 antibody detected the SNX10-HA immunoprecipitated by the anti-HA beads, indicating that the expected SNX10-HA protein was produced. The cells were induced in a similar manner with RANKL, indicated by the increased expression of CLC-7. The anti-SNX10 antibody detected WT SNX10 at 25kDa in the WT cells (asterisk), but only the SNX10-HA protein in the SNX10-HA clone (double asterisk). This indicates that the SNX10-HA cells produce only this SNX10 protein product.

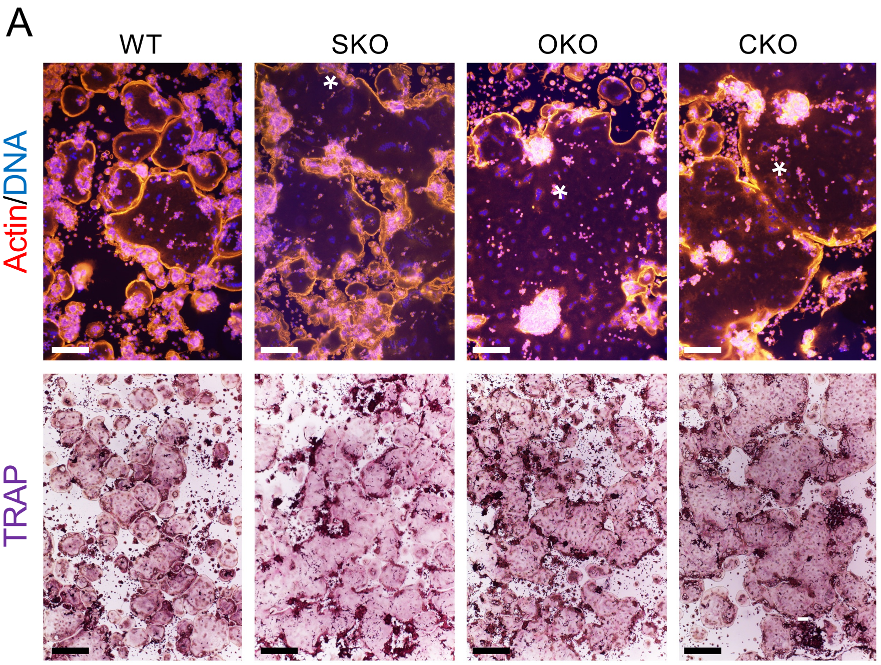


**Supplementary Figure 3 (Identical to Figure 1A, but without indicaiton of cell limits): OCLs lacking SNX10 (SKO), OSTM1 (OKO), or CLCN7 (CKO) are gigantic. A.** RAW 264.7 cells of the indicated genotypes were plated on glass coverslips and induced to differentiate with M-CSF and RANKL. Cells were stained for actin (red/orange) and DNA (blue) (top) or tartrate-resistant acid phosphatase (TRAP) (bottom). Each of the mutant images contains part of a single cell (marked by an asterisk) that extends significantly beyond the field of view. Scale bars: 200 mm (top), 500 mm (bottom). WT- control, non-targeted cells.

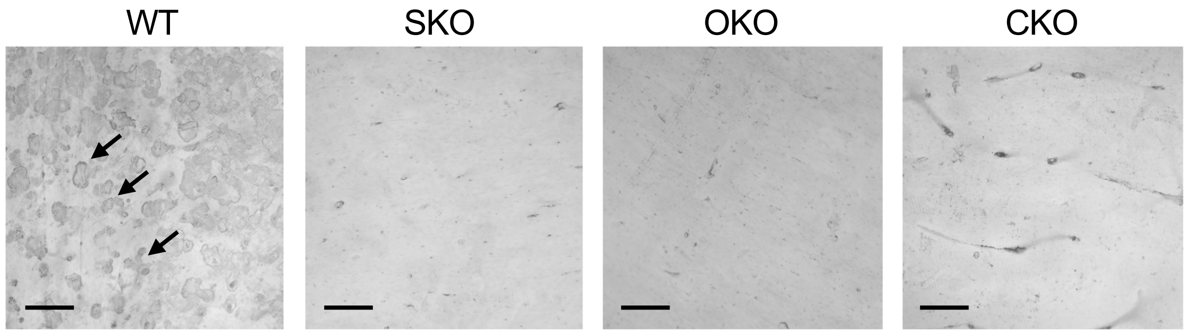


**Supplementary Figure 4 OCLs from homozygous SKO, OKO and CKO mice cannot degrade bone.** Spleen cells from mice of the indicated genotypes were seeded on bone and differentiated into OCLs. Cells were then removed, and the bone surface was stained with WGA/DAB. Resorption pits were observed only in the WT control samples (e.g., arrows). Scale bars: 200 mm.

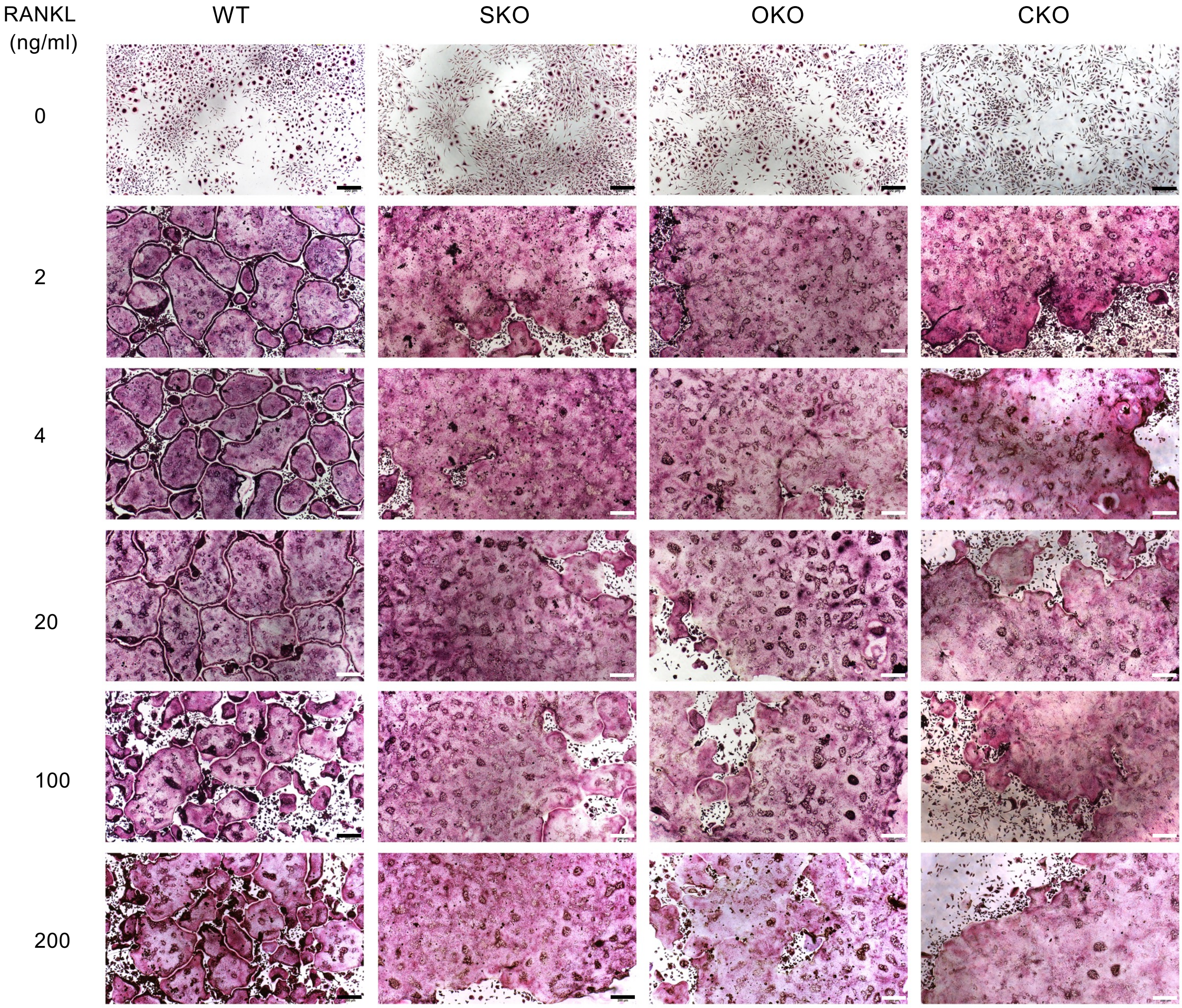


**Supplementary Figure 5: Regulated vs. deregulated OCL fusion is independent of RANKL concentration**. Equal numbers of M-CSF-treated splenocytes were cultured with 20 ng/ml M-CSF and the indicated concentrations of RANKL for 3-6 days and then stained for TRAP. Each WT image contains multiple juxtaposed OCLs, while each mutant image contains part of a single OCL. Scale bars: 200 μm.

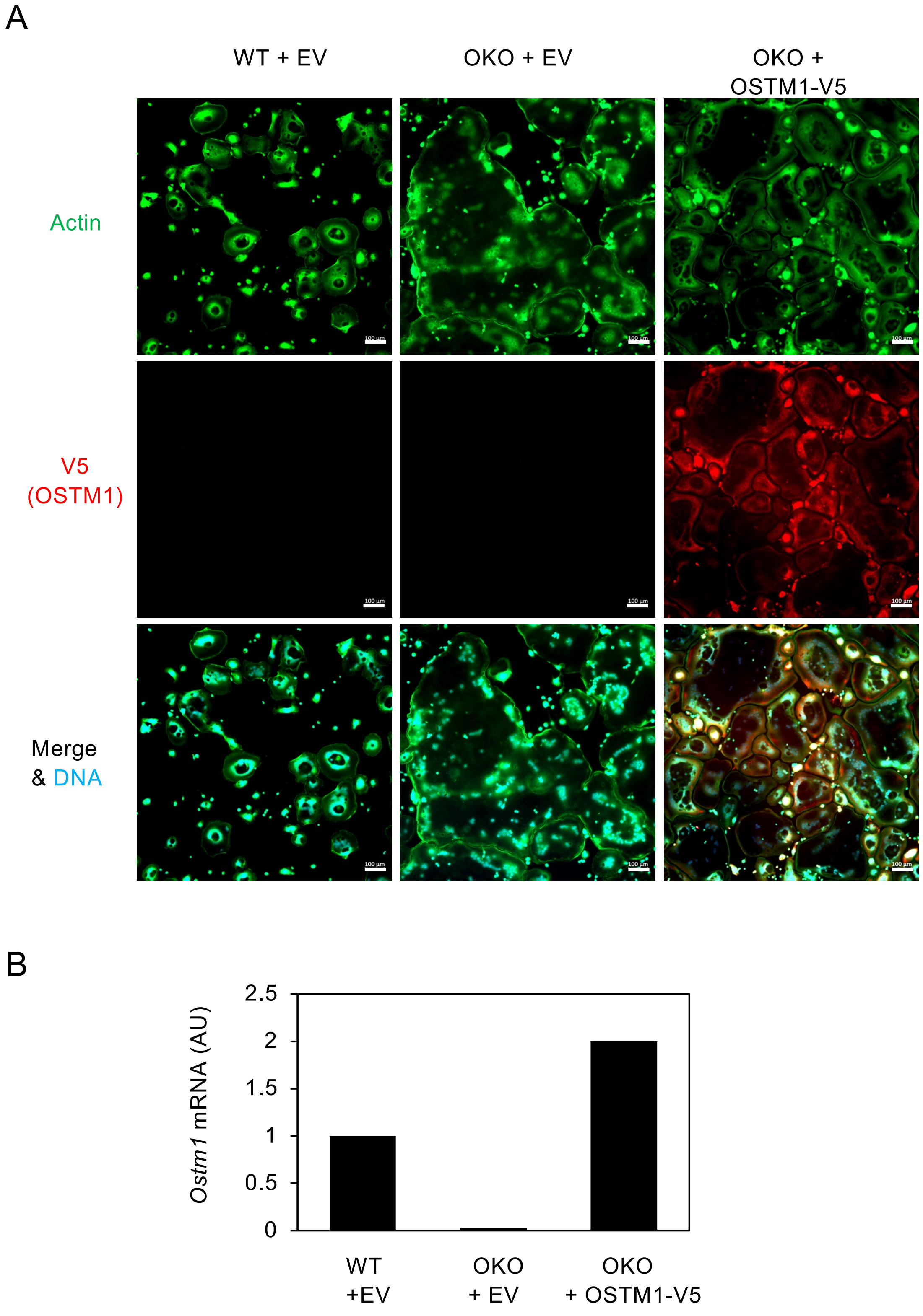


**Supplementary Figure 6: Exogenous OSTM1 rescues the hyperfusion phenotype of OKO OCLs. A.** WT or OKO RAW264.7 cells were infected with lentiviruses for expression of V5-tagged OSTM1 or empty vectors (EV). Scale bars: 100 µm. **B**. qPCR of *Ostm1* mRNA in the indicated cells. Shown is a representative experiment of three performed.

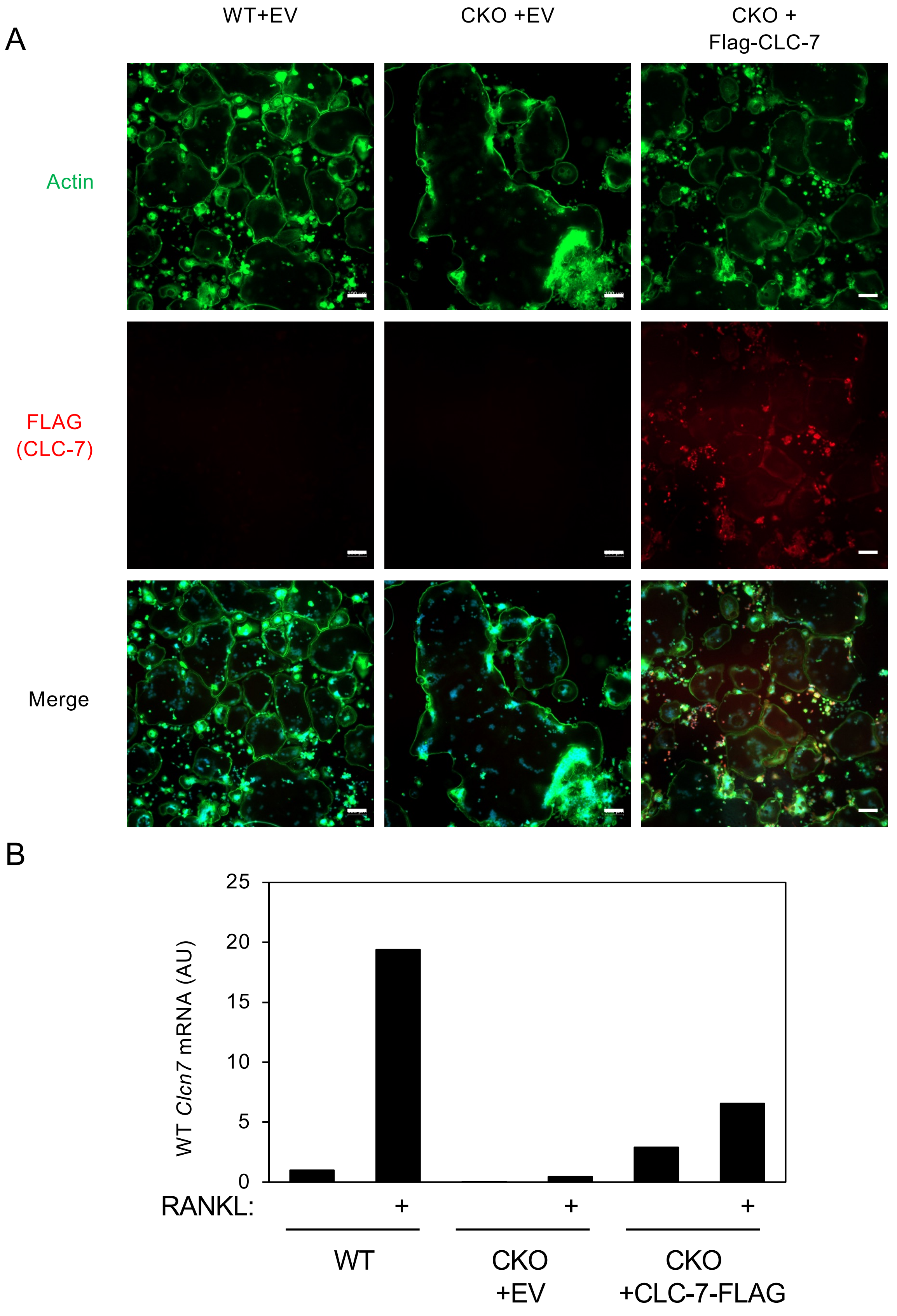


**Supplementary Figure 7: Exogenous CLC-7 rescues the hyperfusion phenotype of CKO OCLs. A.** WT or CKO RAW264.7 cells were infected with lentiviruses for expression of FLAG-tagged CLC-7 or empty vector (EV). Scale bars: 100 µm. **B.** qPCR of *Clcn7* mRNA in the indicated cells. Note that the primer pair used (*Clcn7* set g, Supplemental Table 2) recognizes only the WT mRNA that produces WT CLC-7 protein. This primer pair does not recognize mRNA produced from the endogenous targeted allele in CKO cells, which does not produce protein (Supplemental Figure 1).

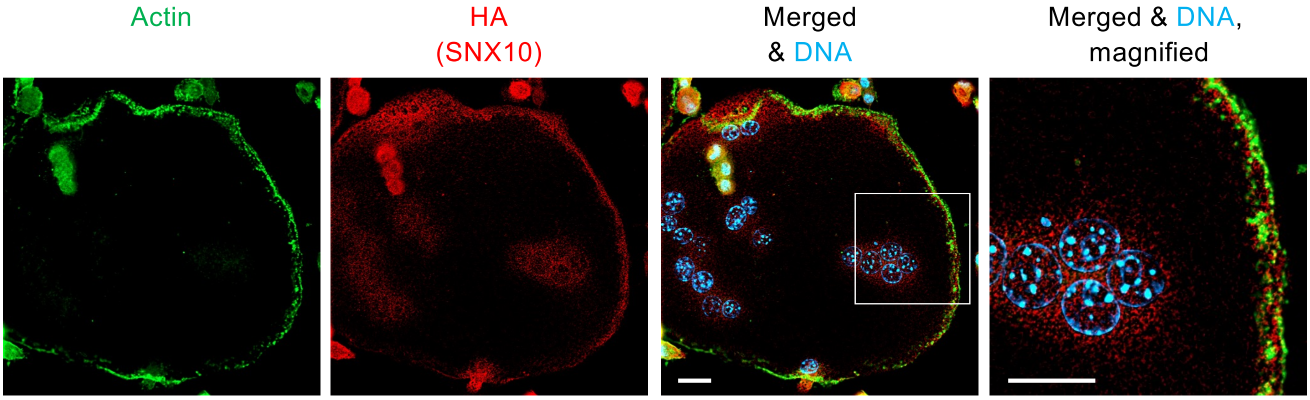


**Supplementary Figure 8 SNX10 that is located at the cell periphery does not overlap with the podosomal ring in OCLs.** WT RAW264.7 OCLs were stained for actin (phalloidin) and for SNX10 (via the C-terminal HA tag inserted into the C terminus of the endogenous protein). Merged image includes also staining for DNA (blue). Scale bars: 20 µm.

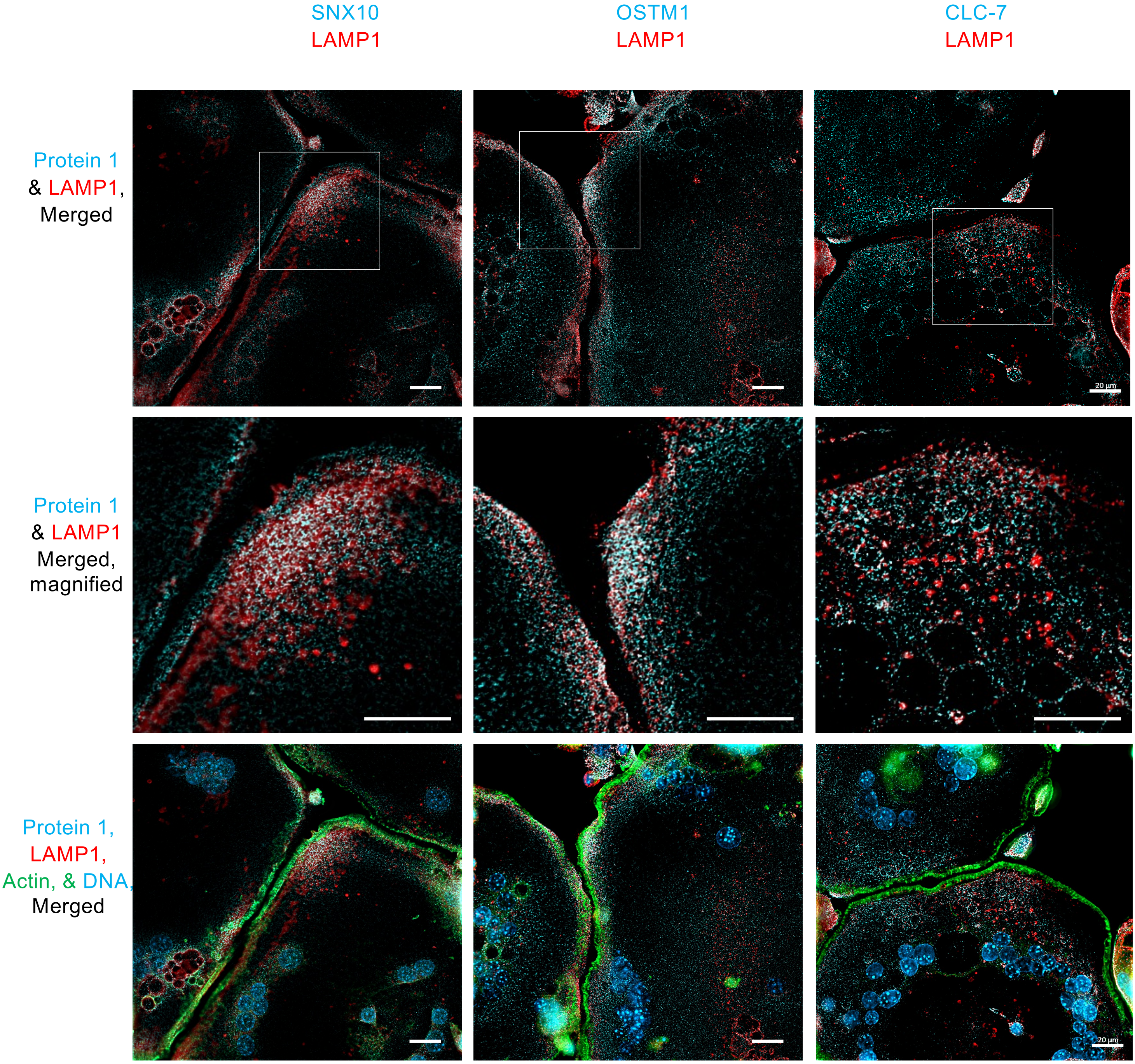


**Suplementary Figure 9: Simiar to Figure 6, Co-localization of SNX10, OSTM1, or CLC-7 with LAMP1 in OCLs,** showing overlap (white) in a broader enlarged region. RAW264.7 cells were differentiated into OCLs and stained as indicated. As described in the Results, endogenous SNX10 was visualized via a C-terminal HA tag that was inserted into the *Snx10* gene; exogenous, V5-tagged OSTM1 was expressed in OSTM1-KO cells, and exogenous FLAG-tagged CLC-7 was expressed in CLC-7-KO cells. Scale bars: 20 μm.

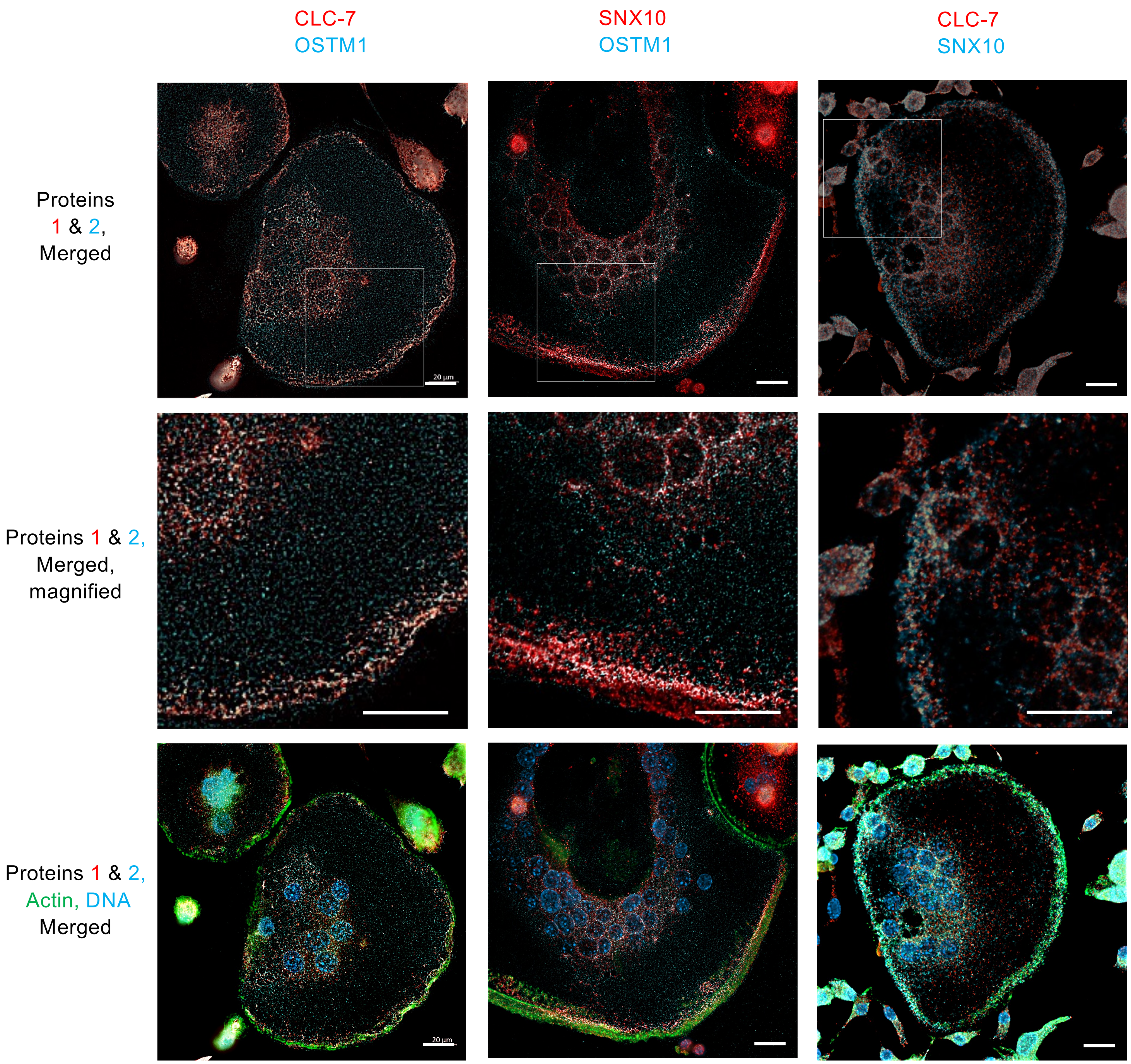


**Supplementary Figure 10: Similar to Figure 7, Co-localization of SNX10, OSTM1, and CLC-7 in RAW264.7 OCLs,** showing overlap (white) in a broader enlarged region. Endogenous SNX10 was visualized via a C-terminal HA tag, exogenous OSTM1 was visualized via a C-terminal V5 tag, and endogenous CLC-7 was visualized using anti-CLC-7 antibodies. For SNX10-OSTM1 co-localization, cells expressing SNX10-HA were co-cultured with cells expressing OSTM1-V5; cell fusion produced cells that express both proteins for analysis. Scale bars: 20 µm.
